# The phenotypic landscape of essential human genes

**DOI:** 10.1101/2021.11.28.470116

**Authors:** Luke Funk, Kuan-Chung Su, David Feldman, Avtar Singh, Brittania Moodie, Paul C. Blainey, Iain M. Cheeseman

**Affiliations:** Broad Institute of MIT and Harvard; 415 Main St., Cambridge, MA 02142, USA; Harvard-MIT Health Sciences and Technology, Massachusetts Institute of Technology, Cambridge, MA 02142, USA; Whitehead Institute for Biomedical Research; 455 Main Street, Cambridge, MA 02142; Department of Biology, Massachusetts Institute of Technology, Cambridge, MA 02142; Department of Biological Engineering, Massachusetts Institute of Technology, Cambridge, MA 02142; Koch Institute for Integrative Cancer Research at MIT, Cambridge, MA 02142

## Abstract

Understanding the basis for cellular growth, proliferation, and function requires determining the contributions of essential genes to diverse cellular processes. Here, we combined pooled CRISPR/Cas9-based functional screening of 5,072 fitness-conferring genes in human cells with microscopy-based visualization of DNA, DNA damage, actin, and microtubules. Analysis of >31 million individual cells revealed measurable phenotypes for >90% of genes. Using multi-dimensional clustering based on hundreds of quantitative phenotypic parameters, we identified co-functional genes across diverse cellular activities, revealing novel gene functions and associations. Pooled live-cell screening of ∼450,000 cell division events for 239 genes further identified functional contributions to chromosome segregation. Our work creates a resource for the phenotypic analysis of core cellular processes and defines the functional landscape of essential human genes.

---

For a human cell to grow, proliferate, and function, it must carry out a variety of essential cellular processes, including transcription, mRNA splicing, translation, vesicle trafficking, proteolysis, DNA replication, and cell division. CRISPR/ Cas9-based pooled genetic screens have revolutionized the ability to test the functional requirements for cell growth and proliferation by enabling the potent disruption of thousands of individual genetic elements in single experiments (*1*). However, most current screening approaches, including those based on fluorescence-activated cell sorting (FACS) of cell populations (*2, 3*), produce a single scalar measurement of barcode enrichment or depletion that summarizes the contributions of each perturbation to cellular phenotypes at the population level. In cellular fitness screens using these approaches, it is thus rarely possible to distinguish between essential genes that function in distinct cellular processes. To improve the differentiation of complex phenotypes, recent studies have combined pooled functional genetic screens with additional measurements at limited scales, including single-cell profiling of transcriptional cell states (*4*). Defining the specific contributions of essential genes to core cellular processes requires a quantitative analysis of complex cellular phenotypes, many of which can be directly visualized using microscopy. Leveraging the power of microscopy, recent work utilized targeted fluorophore photoactivation of cells exhibiting specific optical phenotypes to enable visual CRISPR screens (*5*–*7*). However, these approaches similarly produce a single enrichment score for each gene, with one predefined phenotype at a time. The ability to interrogate and systematically compare a large and diverse array of cell biological phenotypes simultaneously across thousands of genomic perturbations represents an important unmet goal for functional studies. Here we use optical pooled screening (*8, 9*) to combine large-scale Cas9-based targeting of essential genes with microscopy and image-based profiling of single-cell resolved cell biological phenotypes at a large scale (Fig. 1A).

**Fig. 1.**
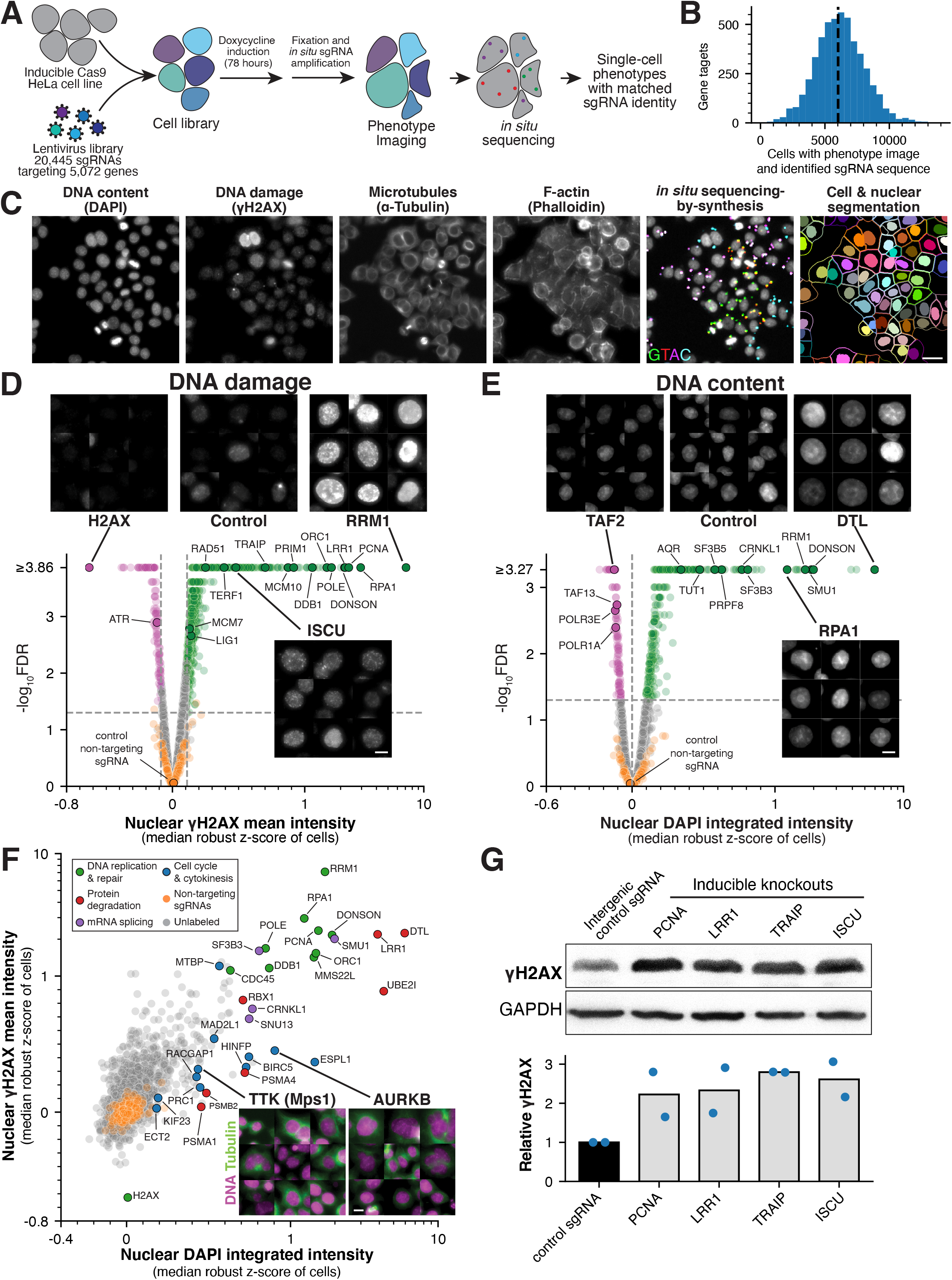
Large-scale image-based pooled CRISPR screen identifies essential genes with roles in genome integrity. **(A)** Experimental workflow for the fixed-cell, image-based pooled CRISPR screen (also see Methods). **(B)** Histogram showing the number of cells analyzed for each gene target with an acquired phenotype image and single sgRNA sequence mapped in situ. **(C)** Example image from the pooled screen showing the phenotype stain channels (DNA, γH2AX, tubulin, and F-actin) together with a matched field-of-view of fluorescent in situ sequencing (Laplacian-of-Gaussian filtered) and cell segmentation. Scale bar, 25 µm. **(D)** Volcano plot for mean nuclear γH2AX intensity across gene targets in the screen. Selected images from the screen show γH2AX staining for example cells to highlight specific targets whose knockout results in increased (green) or decreased (magenta) DNA damage relative to non-targeting control sgRNAs (orange; FDR<0.05). The median robust z-score was measured across cells with the same sgRNA, and aggregated to the gene level by taking the median of sgRNAs targeting the same gene (see Methods). The median robust z-score is plotted on a symmetric log scale (linear between -1 and 1). Raw P-values were computed by comparing gene targets to a bootstrapped null distribution of cells expressing non-targeting sgRNAs (see Methods), with false discovery rate (FDR) estimated using the Benjamini-Hochberg procedure. Scale bar, 10 µm. **(E)** Volcano plot as in (D) for integrated nuclear DAPI intensity, along with example images of DAPI staining for gene knockouts that result in increased or decreased DNA content. Scale bar, 10 µm. **(F)** Scatter plot comparing the relationship between DNA damage and DNA content. A subset of gene knockouts result in particularly increased DNA content due to cell division failure; labeled genes are colored by functional category. Example images show tubulin (green) and DNA (magenta) to highlight polyploid and multinucleate cells. Scale bar, 10 µm. **(G)** Western blot (top) and quantification (bottom) confirming the presence of increased DNA damage based on cellular γH2AX levels for genes identified in the image-based pooled screen and PCNA as a positive control. Each sample represents a distinct cell line with an inducible Cas9 and stably-expressed sgRNA targeting either the indicated gene or a negative control single copy locus. Blue dots indicate independent γH2AX quantifications referenced to GAPDH and relative to the negative control.

## A large-scale, image-based pooled CRISPR screen of essential genes

To determine the functional contributions of essential genes in cultured human cells, we first defined a set of fitness-conferring genes based on combined evidence from multiple Cas9- and transposon-based genetic screens (*9*–*17*; Fig. S1A-B; Methods). This approach defined a collection of 5,072 genes that contribute to optimal cellular fitness, although we note that not every gene will be required for fitness in a given cell line. To create a library of CRISPR sgRNAs targeting this gene collection, we selected four sgRNA sequences per target gene from optimized sgRNA libraries (*18*–*20*), prioritizing guides with evidence of high on-target efficiency and low off-target activity (Methods). In addition, we selected 250 “non-targeting” sgRNAs that lack targets in the human genome as negative controls. Together, this constituted a library of 20,445 total sgRNAs. We delivered the sgRNA library to HeLa cells containing an integrated, doxycycline-inducible Cas9 construct (*21*) using the CROPseq-puro-v2 lentiviral vector that has an optimized sgRNA scaffold (*8, 21, 22*; Methods). Based on a trial image-based screen targeting 400 genes (Fig. S1C) and an analysis of sgRNA depletion from our library at 3 and 5 days post-Cas9 induction (Fig. S1D), we defined a time point at 78 hours post-Cas9 induction to maximize observable phenotypes, balancing the time required for protein depletion with negative fitness effects that cause knockout cells to drop out of the population. For the primary image-based pooled screen, we fixed the cell population at 78 hours post-Cas9 induction and amplified the sgRNA sequences *in situ* as described previously (Fig. 1A, B; *8*). Following amplification, we stained and imaged cells for DNA (DAPI), DNA damage (γH2AX; anti-phospho-Ser139 H2AX antibody), microtubules (anti-α-tubulin antibody), and filamentous actin (phalloidin) (Fig. 1C). These stains were chosen to visualize diverse cell biological behaviors, including nuclear morphology, cell size, DNA damage response, cytoskeletal structures, cell cycle stage, and mitotic chromosome alignment.

Following the completion of phenotype imaging, we performed *in situ* sequencing-by-synthesis to identify the sgRNA present in each individual cell (Fig. 1A, C; Fig. S1E) (*8*), allowing us to directly assess the phenotypic consequences of disrupting each target gene. We extracted 1,084 phenotypic parameters from each individual cell image, including measurements of the intensity and subcellular distribution of each stain, colocalization of stains, and cellular and nuclear size and shape (Methods; Data S1). We identified interphase and mitotic cells as separable cellular states in our dataset with distinct baseline phenotypes. Thus, for phenotype analysis, we classified mitotic and interphase cells using a support vector classifier with a subset of extracted phenotype parameters and conducted downstream analyses separately (Methods; Fig. S1F). Together, this approach yielded microscopy images, extracted phenotypic measurements, and matched sgRNA identities for 31,884,270 individual cells with a median of 6,119 cells per gene target across each set of four sgRNAs (Fig. 1B; Data S1). Image montages and phenotypic parameters of interphase and mitotic cells are available for exploration through the companion interactive web portal (https://nematode.wi.mit.edu/vesuvius/).

## Interphase nuclear phenotypes reveal established and novel regulators of genomic integrity

Maintaining genomic integrity is critical to ensuring proper cellular function, as DNA mutations and chromosome imbalances result in genome instability, misregulated gene expression, cell inviability, and oncogenic cell states. Cells utilize a range of DNA damage-sensors, DNA repair mechanisms, and cell cycle checkpoints to recognize and correct genomic aberrations and protect the genome against spontaneous DNA damage, DNA replication-induced errors, and chromosome segregation defects (*24*). To identify genes that are required for genome integrity, we analyzed nuclear phenotypic parameters in interphase cells from our screen that monitor DNA damage (mean γH2AX nuclear intensity) and total DNA content (integrated DAPI nuclear intensity; Fig. 1D, E). We defined summary phenotype scores for each gene target as the median robust z-score of cells relative to the local intermixed population expressing non-targeting control sgRNAs (Methods). Gene targets that displayed decreased γH2AX intensity in interphase cells included H2AX itself and ATR, which is involved in directing the γH2AX phosphorylation event (Fig. 1D). Reciprocally, of the 5,072 genes targeted in the screen, we observed 1,693 genes whose disruption resulted in significantly increased γH2AX intensity (Methods). The top scoring hits included many factors with known roles in DNA replication (e.g., RRM1, PCNA, DNA Polymerase subunits, Primase subunit 1, DNA Ligase 1, RPA1/2/3, ORC and MCM2-7 subunits), DNA repair (e.g., RAD51, REV3L, ATAD5, DONSON, DTL, DDB1), and telomere protection (TERF1/2, RTEL1) (Fig. 1D). Many gene targets that caused increased DNA damage also resulted in increased DNA content (Fig. 1E), including most of the knockouts of DNA replication and repair factors listed above. Gene knockouts with increased γH2AX and DNA content were also enriched for spliceosome components (Fig. S2A-B), consistent with reports that disrupting mRNA splicing results in a DNA damage response (*25*). We observed an overall correlation between DNA damage and total DNA content (r = 0.62), although some knockouts displayed strong increases in DNA content but less severe DNA damage (Fig. 1F). This includes many proteasome 20S core particle subunits and gene targets whose disruption prevents cytokinesis (AURKB, BIRC5, CDCA8, PRC1, KIF23, ECT2) or that allow cells to progress through cell division without segregating their chromosomes (ESPL1, TTK/Mps1, MAD2L1). Targeting each of these cytokinesis and chromosome segregation genes results in more cells with increased DNA content and nuclear area due to tetraploidy or multinucleation (Fig. 1F; Fig. S2C-E).

In addition to the established players in DNA replication and repair, we identified multiple gene targets with poorly established functional roles whose knockouts resulted in increased DNA damage. We noted that a substantial portion of these gene targets are present as multi-gene clusters within a single chromosome and therefore likely exhibit irreparable DNA damage as a result of sgRNAs targeting multiple loci (Fig. S2F). We also identified increased DNA damage following knockout of the E3 ubiquitin ligase subunits LRR1 and TRAIP, and the mitochondrial iron-sulfur cluster biogenesis gene ISCU. To confirm their roles in DNA damage, we generated cell lines with inducible Cas9 expression and a single sgRNA targeting the corresponding genes (*21*). Based on Western blotting, we noted a substantial increase in γH2AX levels following ISCU, LRR1, and TRAIP depletion, along with PCNA as a positive control, compared to a control sgRNA with a single target site (Fig. 1G). The effect of ISCU knockout is consistent with the requirement for iron-sulfur clusters in the enzymatic activity of proteins involved in DNA metabolism (*26*), and LRR1 and TRAIP have recently been reported to play roles in replisome disassembly (*27*). Together, this analysis highlights the importance of multiple genes in DNA replication and repair, and validates our image-based screening strategy to identify diverse players in genome integrity.

## Identification of essential genes controlling cytoskeletal function

To direct cellular proliferation, structure, organization, and mechanical force production, cells rely on complex and dynamic cytoskeletal networks involving actin and microtubule polymers (*28, 29*). Our screen measured interphase cytoplasmic features for both filamentous-actin (F-actin) and microtubules, enabling the broad identification of essential genes involved in cytoskeletal processes and cellular organization. An analysis of interphase mean F-actin intensity revealed 460 gene knockouts with decreased intensity relative to non-targeting controls and 899 genes with increased intensity (Fig. 2A; Fig. S3A). Amongst these genes, we identified established factors required for regulating actin assembly and dynamics. For example, knockout of the actin depolymerization and severing factor cofilin (CFL1) or capping protein (CAPZB), which acts to block actin elongation, resulted in substantially increased actin polymer levels. In contrast, depletion of RHOA or ARHGEF7, which regulate the formation of actin fibers, resulted in strongly decreased actin levels. Although the Arp2/3 complex plays an established role in nucleating actin assembly, we found that its loss resulted in increased cellular actin intensity. However, this was coupled with a substantial decrease in interphase cell area (Fig. S3B). This suggests that disrupting selected components of the actin cytoskeleton also perturbs cellular adhesion, resulting in reduced cell-substrate contacts and an increase in mean cytoplasmic actin intensity due to altered cell shape. Indeed, we observed a similar phenotype for the adhesion components RAC1, Integrin subunits (ITGAV, ITGB1, and ITG5), TLN1, CRKL, ILK, and many others (Fig. S3B, C). The gene target whose loss resulted in the largest increase of mean F-actin intensity in both interphase and mitotic cells was the E3 ubiquitin ligase KCTD10, along with its partners CUL3 and RBX1 (Fig. 2A; Fig. S3D). Recent work implicated KCTD10 in restricting actin assembly during cell migration or developmentally-programmed cell fusion (*30, 31*), but our analysis suggests a general role for this E3 ubiquitin ligase in regulating actin assembly.

**Fig. 2.**
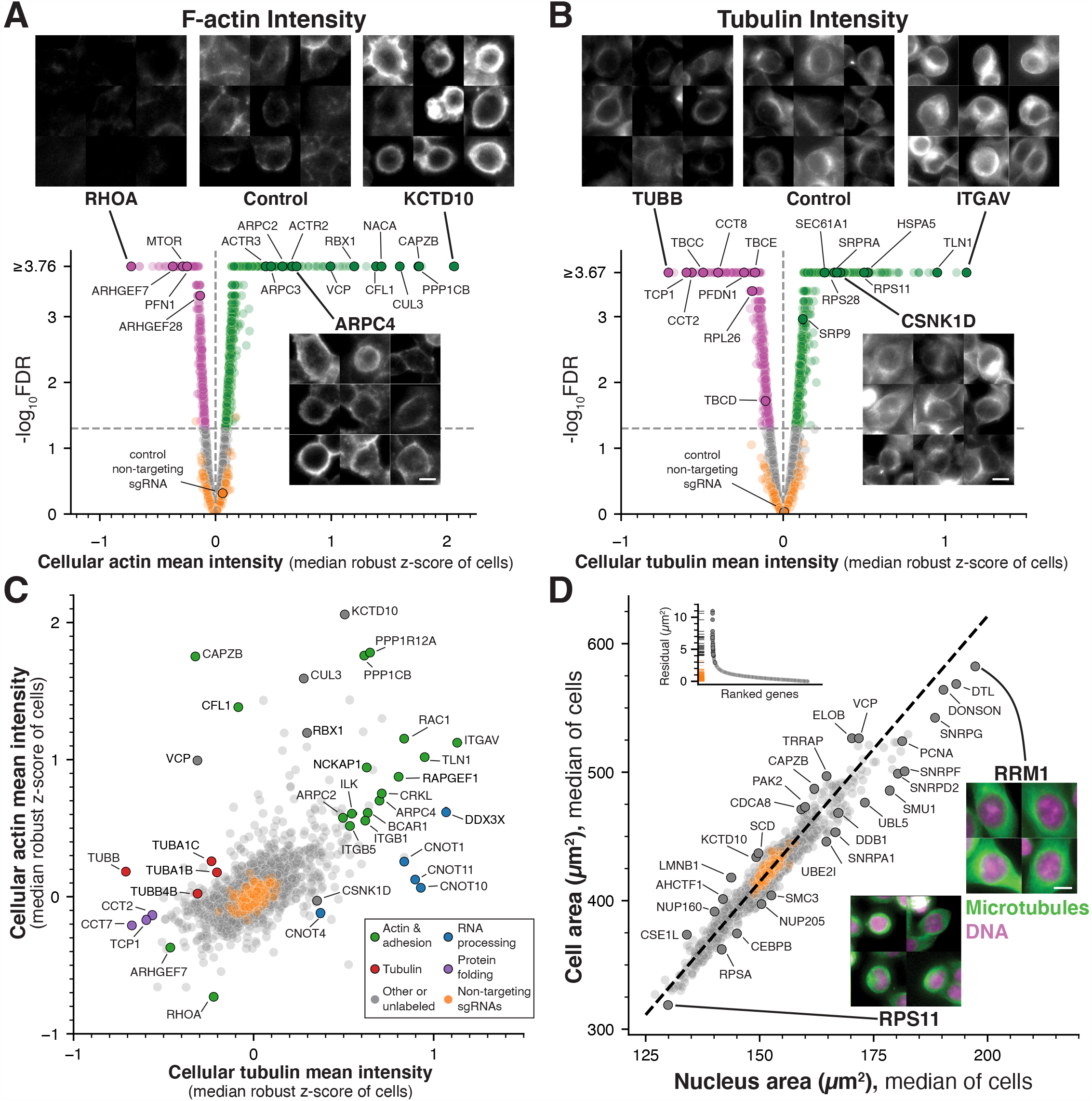
Identification of essential genes regulating cytoskeletal structures and cellular organization. (**A**) Volcano plot for mean cellular F-actin (phalloidin) intensity across gene targets in the screen. Selected images from the screen show phalloidin staining for example cells to highlight specific targets that result in increased (green) or decreased (magenta) cellular actin levels relative to non-targeting control sgRNAs (orange; FDR<0.05). The median robust z-score was measured across cells with the same sgRNA, and aggregated to the gene level by taking the median of sgRNAs targeting the same gene (see Methods). Raw P-values were computed by comparing gene targets to a bootstrapped null distribution of cells expressing non-targeting sgRNAs (see Methods), with false discovery rate (FDR) estimated using the Benjamini-Hochberg procedure. (**B**) Volcano plot as in (A) for mean cellular tubulin intensity, along with example images of tubulin staining for gene knockouts that result in increased or decreased tubulin levels. (**C**) Scatter plot comparing the relationship between actin and tubulin stain intensity highlights gene targets that selectively affect one cytoskeletal element. A subset of knockouts impact both actin and tubulin mean intensity measurements by disrupting cellular adhesion and decreasing the segmented cell area (see also Fig. S3B, C, F). Labeled genes are colored by functional category. (**D**) Scatter plot showing the comparison between cellular and nuclear area across gene targets. These morphological features are highly correlated across scales and conditions (r = 0.96). The median of area measurements was computed across cells with the same sgRNA and aggregated to the gene level by taking the median of sgRNAs targeting the same gene. Orthogonal regression was performed to identify gene targets that result in an altered nuclear:cytoplasmic area ratio (dotted line). Labeled genes are also highlighted in the distribution of regression residuals (inset). Example images display DNA (magenta) and tubulin (green) staining for gene targets that result in increased or decreased cell and nuclear size. Scale bars, 10 µm.

We also identified multiple factors regulating interphase tubulin levels. Mean tubulin intensity was significantly decreased for 492 gene targets when compared to non-targeting controls, including genes encoding tubulin proteins (TUBA1B/C, TUBB, TUBB4B), tubulin-specific chaperones (TBCC/D/E), and factors required for tubulin folding and complex assembly (CCT chaperonins/TRiC complex and prefoldin subunits; Fig. 2B). Reciprocally, we observed an increased mean tubulin intensity for 639 knockouts. However, as noted above, cytoplasmic proteins such as actin and tubulin may display increased mean stain intensity under conditions where cell area is reduced due to altered substrate adhesion (Fig. S3B, C, F). Thus, we compared actin and tubulin intensity to identify gene targets that selectively affect one stain (Fig. 2C). We observed substantially increased tubulin fluorescence, but not increased actin intensity, for Casein kinase I delta (CSNK1D), which has been suggested to regulate microtubule-associated proteins (*32*), and subunits of the CCR4-NOT complex (CNOT1/4/10/11), which functions in post-transcriptional mRNA regulation (*33*). Together, this analysis of interphase cytoplasmic actin and tubulin intensity reveals the contributions of diverse molecular players for roles in controlling cytoskeletal assembly and dynamics.

## Analysis of morphological phenotypes reveals a tight correspondence between cellular and nuclear size

In addition to measuring stain intensity for each marker, we also measured multiple morphological parameters including nuclear and cellular area. We noted substantial differences in median interphase cell area across the different gene targets, ranging in segmented area from 319 µm^2^ to 583 µm^2^ (Fig. 2D; Fig. S3G). Consistent with a required role for protein production in cell growth, targeting ribosome and ribosome biogenesis genes resulted in substantially reduced cell area (Fig. S3H). In contrast, gene targets with roles in DNA replication and repair, mRNA splicing, and proteasome function displayed increased cell areas (Fig. S3H), suggesting continued cell growth in the absence of further division. Strikingly, we observed a strong correlation between cell area and nuclear area across all tested gene targets (r = 0.96; Fig. 2D). Prior work has suggested that cells actively regulate the size ratio between their nucleus and cytoplasm (*34*). Our analysis demonstrates that, across a wide range of cell sizes and functional perturbations, this relationship is closely maintained. Although we observed a clear relationship between nuclear size and cell size, we identified a limited number of gene targets whose depletion disrupted this coordinated scaling (Fig. 2D). A subset of gene targets displayed abnormally large nuclei for their given cell size, including RNA splicing factors (e.g., SMU1 and UBL5), the nuclear pore complex member NUP205, and DNA replication factors (e.g, PCNA, DDB1, RRM1). We also identified gene knockouts that displayed decreased relative nuclear size, including lamin B1 (LMNB1), the nucleocytoplasmic transport protein CSE1L, and the nuclear pore components NUP160 and AHCTF1, consistent with roles in controlling nuclear integrity and function. Together, this analysis demonstrates that cell biological parameters from a large-scale screen can be used to reveal broader paradigms for the control of cell size and organization.

## Phenotypic clustering of interphase cellular parameters defines co-functional genes

We next sought to take advantage of the full range of identifiable phenotypes in the rich image data from our screen to reveal additional gene activities required for cellular function. To define the phenotypic landscape of essential genes, we selected and aggregated 472 non-redundant parameters extracted from each interphase cell image to create a summary phenotypic profile for each gene target. We then analyzed these profiles in a high-dimensional space using the PHATE algorithm (*35*) and performed clustering to identify genes with similar phenotypes (Methods; Fig. 3A; Fig. S4A-C). Based on a comparison of knockout phenotypes to non-targeting guides, 4,665 of the 5,072 gene targets in our screen display a measurable interphase phenotype (Fig. S4A-B). Of the remaining 407 gene knockouts, only 55 genes displayed strong fitness effects at 5 days post-Cas9 induction based on sgRNA depletion from our library (Fig. S4B). Thus, the 352 genes without a measurable phenotype or fitness effect are likely not required for cellular fitness in HeLa cells at the tested time point following Cas9 induction.

**Fig. 3.**
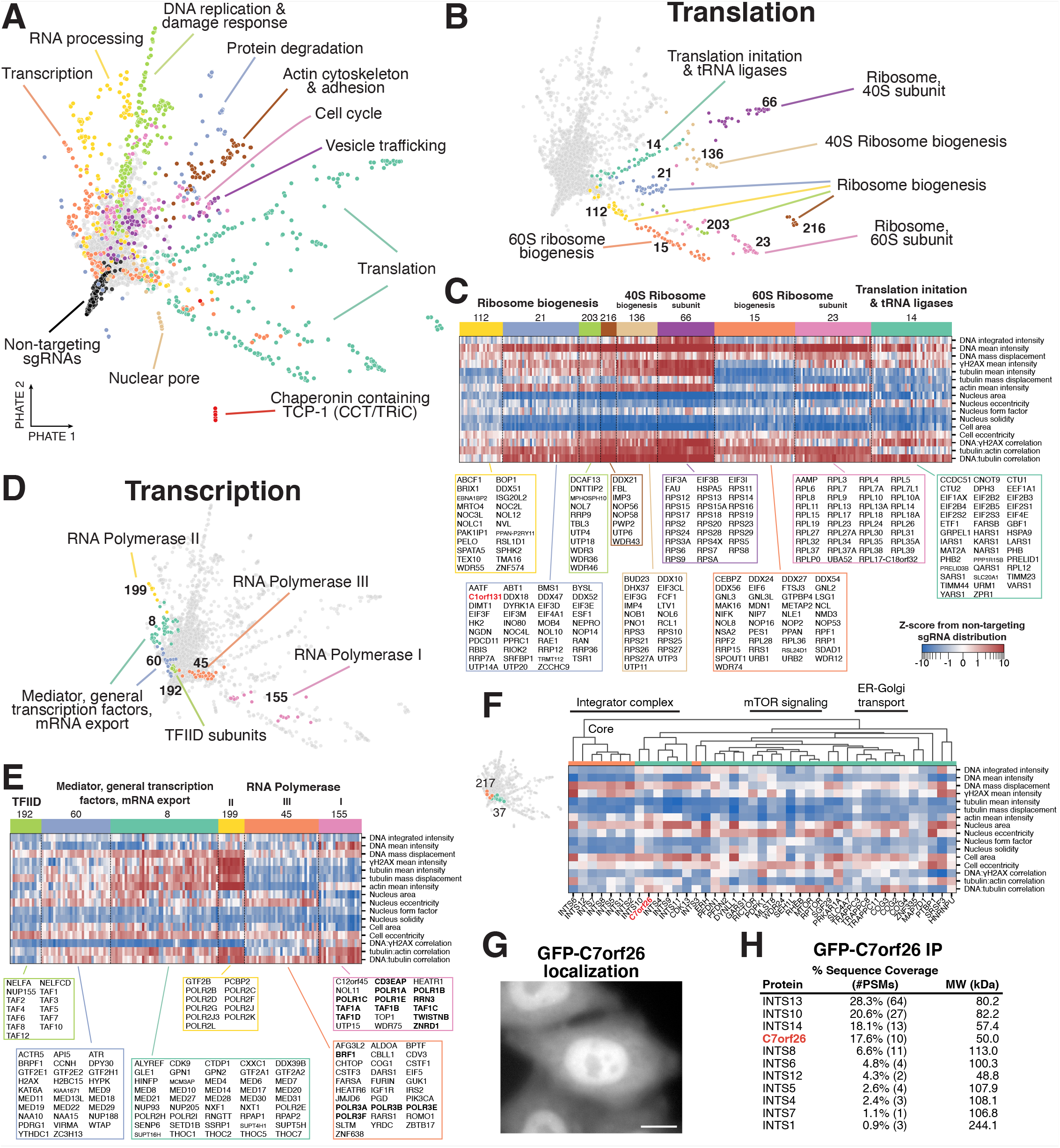
Clustering of multi-dimensional interphase phenotypes reveals co-functional essential genes. (**A**) Two-dimensional representation of the interphase phenotype landscape of gene targets in the primary screen computed using PHATE (*35*) with hundreds of summary phenotype parameters, and then clustered to form groups of genes with similar phenotypes (see Methods). Each dot represents a single gene target, colored corresponding to the indicated functional category of grouped clusters. (**B**) Individual clusters of genes related to translation from (A) identify fine-grained sub-categories of gene function in ribosome biogenesis, translation initiation, and individual ribosome subunits. Functional descriptions of labeled cluster numbers summarize the roles of the contained gene targets. (**C**) Heat map of interphase knockout phenotypes corresponding to the translation clusters in (B) for a manually-selected subset of phenotype parameters. This highlights the phenotypic similarity of gene targets within functionally-coherent clusters, but clear distinctions between separate clusters despite broadly related roles in translation. All genes from each cluster are listed below. Parameters are presented as z-scores from the distribution of non-targeting sgRNAs, visualized on a symmetric log scale (linear between -1 and 1). (**D**) Individual clusters of genes related to transcription from (A) indicate separate clusters for components of each type of RNA polymerase, TFIID, and related complexes. (**E**) Heat map as in (C) corresponding to the clusters in (D) highlighting the phenotypic similarities and differences that define each cluster of genes with transcriptional functions. (**F**) Heat map as in (C) of interphase clusters 37 and 217, demonstrating the phenotypic similarity between C7orf26 knockouts with those of the integrator complex. Hierarchical clustering (top) within these clusters using the correlation of high-dimensional phenotype profiles (see Methods) indicates particularly strong similarities between C7orf26 (red) and INTS10, and predicts that the uncharacterized gene C7orf26 is co-functional with established Integrator subunits. Hierarchical clustering within cluster 37 also implicates an association between mTOR signaling components and ER-Golgi transport factors. (**G**) Fluorescent image of human cells expressing GFP-C7orf26 demonstrates nuclear localization. Scale bar, 10 µm. (**H**) Mass spectrometry from an immunoprecipitation of GFP-C7orf26 in human cells relative to controls indicates that C7orf26 associates with subunits of the Integrator Complex.

From the PHATE clustering analysis, we noted clear functional relationships between the genes within a given cluster, allowing us to identify major clusters primarily composed of gene targets with roles in transcription, RNA processing, translation, protein degradation, DNA replication and damage response, cell cycle control, or other core cellular processes (Fig. 3A-E; Fig. S5; https://nematode.wi.mit.edu/vesuvius/). Strikingly, the clustering behaviors also allowed us to distinguish high-resolution functional sub-categories within each cellular process. For example, despite a shared role in translation, we identified separable clusters containing established 40S ribosome subunits (cluster 66), 60S ribosome subunits (cluster 23), tRNA ligases and eIF2 translation initiation subunits (cluster 14), distinct clusters for factors involved in 40S ribosome biogenesis (cluster 136) and 60S ribosome biogenesis (cluster 15), and several others which included nucleolar proteins, RNA helicases, and additional factors involved in translation initiation or ribosome biogenesis (clusters 21, 112, 203, and 216; Fig. 3B-C). Knockouts for the genes within each of these clusters resulted in reduced nuclear and cellular areas, but displayed differences in other cellular phenotypic parameters, such as actin and tubulin staining intensities, enabling distinction among these functional sub-categories (Fig. 3C). Similarly, we identified multiple clusters containing 26S proteasome subunits with phenotypic differences that allowed segregation of 20S core particle subunits (167) from the 19S regulatory particle ATPase (106) and non-ATPase components (213), as well as a cluster containing components of the COP9 Signalosome (200), which controls ubiquitin-dependent processes (Fig. S5A). Finally, we observed multiple distinct clusters for the core transcriptional machinery, including TFIID subunits (192), RNA Polymerase II subunits (199), two clusters comprised of Mediator complex subunits, General Transcription Factors (GTFs), and mRNA export factors (8 and 60), and separate clusters containing RNA Polymerase I (155) and RNA Polymerase III (45) components (Fig. 3D-E). Interestingly, although our cellular imaging did not include membrane-targeted markers for cellular organelles, such as the Golgi or Endoplasmic Reticulum, we identified multiple distinct clusters comprised of vesicle trafficking components. For example, we identified a cluster (201) containing the coatamer subunits (COPA/B1/B2/G1/Z1, ARCN1), SNAP proteins (NAPA and GOSR2), and cholesterol biosynthesis proteins HMGCS1 and HMGCR, a second cluster (54) containing signal recognition particle (SRP19/54), ESCRT proteins (UBAP1, CHMP6, and VPS28), clathrin, and additional vesicle trafficking proteins, as well as a cluster (140) containing the exocyst complex (EXOC1/4/5/7) and glycosylation machinery (Fig. S5E). This suggests that specific cell morphological changes resulting indirectly from perturbing vesicle trafficking and organelle function can be detected by our extracted image parameters, beyond what is easily distinguishable by visual inspection.

Together, our work demonstrates that phenotypic clustering using quantitative parameters extracted from cell images provides a fine-grained picture of the distinct functional contributions of specific protein sub-complexes to core cellular processes.

## Phenotypic clustering provides novel insights into gene functions and pathway relationships

The coherent phenotypic similarity of co-functional gene targets within a given cluster is readily apparent based on the numerous observed clusters containing closely related, well-characterized genes. Based on this behavior, the presence of additional genes within a phenotypic cluster provides a powerful prediction for their contributions to cellular function. Analysis of these clusters revealed unexpected connections between cellular processes. For example, cluster 37 contains the Conserved Oligomeric Golgi complex (COG2/3/4) and Trafficking Protein Particle complex (TRAPPC3/8/11), together with established mTOR signaling components including the mTOR complex 1 (MTOR, RPTOR, MLST8), RICTOR, RHEB, and GATOR2 complex components (SEH1L, WDR24; Fig. 3F). Although the regulation of mTOR signaling has focused on its association with the lysosome, this phenotypic clustering supports evidence of Golgi-localized components playing a functional role in mTORC1 regulation (*36*), or suggests a role for these Golgi-derived factors in lysosome biogenesis. Similarly, we found that the DICER1 ribonuclease and the microRNA microprocessor subunit DGCR8 clustered with exocyst complex subunits (cluster 140), instead of with other RNA-associated regulatory factors (Fig. S6A). This supports the association of DICER1 with exosomes (*37*), but suggests a cell autonomous role rather than one facilitating cell-cell communication. Clustering also revealed phenotypic similarities between knockouts of the key signaling proteins KRAS and BRAF and multiple mitochondrial components, such as mitochondrial ribosome subunits and proteins involved in mitochondrial respiration (NADH dehydrogenase and Cytochrome) (cluster 149; Fig. S6B). KRAS and BRAF are mutated in a substantial fraction of cancers, although not in the HeLa cells used in this study. This clustering behavior highlights an important role for KRAS and BRAF signaling in maintaining normal metabolic homeostasis. Finally, although several clusters containing transcriptional regulators exhibited related phenotypes, we identified a cluster (121) with a distinct phenotypic profile that contains both of the master regulator Myc and Max transcription factors, along with multiple other transcriptional regulators (FOXN1, ILF3, SP2, ZBTB11, nuclear respiratory factor 2 genes GABPA and GABPB1), chromatin remodeling factors (ZMYND8, H3K36 methyltransferase SETD2), and E3 ubiquitin ligase components (KEAP1, DDA1; Fig. S6C). This suggests that these factors may either be specifically required for Myc expression (for example, see *38*) or work with Myc to promote downstream expression at its target promoters.

Our clustering analysis further implicated poorly characterized gene targets in specific cellular activities. For example, we nominated C1orf131 as a putative nucleolar component involved in ribosome biogenesis based on its membership in cluster 21 (Fig. 3C), a prediction that was recently confirmed by others (*39*). Similarly, AKIRIN2 clustered with the 20S core particle proteasome subunits (cluster 167, Fig. S5A), and was recently described as a required proteasome nuclear import factor (*40*). In addition, HNRNPD was present in cluster 197 together with METTL3 and METTL14 (Fig. S6B), which form the core heterodimer that writes m6A mRNA modifications, consistent with the emerging role of m6A modifications in promoting HNRNPD associations with mRNA (*41*). We also identified the uncharacterized gene C7orf26 in cluster 37 as displaying phenotypes closely related to those observed in knockouts of known subunits of the Integrator complex, an RNA endonuclease involved in RNA processing (*42*) present in clusters 37 and 217 (Fig. 3F). C7orf26 and Integrator complex knockouts resulted in reduced interphase tubulin intensity without corresponding changes in actin intensity. To evaluate this co-clustering relationship, we generated a cell line stably expressing GFP-C7orf26. This GFP-C7orf26 fusion localized to the nucleus (Fig. 3G), consistent with Integrator complex localization (*43*). Using affinity purifications coupled to mass spectrometry, GFP-C7orf26 pull downs specifically isolated multiple Integrator complex subunits, with particularly robust levels of INTS13, INTS10, and INTS14 (Fig. 3H; also see *44*). This is consistent with the strong phenotype similarity in our screen between C7orf26 and INTS10 knockouts (Fig. 3F). INTS10, INTS13, and INTS14 were recently shown to comprise a functional subunit of the Integrator complex that associates the cleavage module with target RNA (*45*), suggesting C7orf26 may interact with this sub-complex. Thus, the phenotypic clustering of this dataset identifies established interacting partners and provides predictive insights to identify novel associations and co-functional players across key cell biological processes with the potential for many additional discoveries.

## Analysis of mitotic phenotypes identifies requirements for proper cell division

We next analyzed the phenotypes observed in mitotic cells for each gene target. In total, 2.6% of the cells visualized in our microscopy-based screen were present in the mitotic phase of the cell cycle (median of 157 mitotic cells per gene target). In the presence of mitotic errors, cells activate the spindle assembly checkpoint and arrest in mitosis (*46*). Therefore, an increased fraction of obtained cell images that are identified as mitotic for a given gene (i.e., mitotic index) can reflect a mitotic disruption (Fig. 4A). We observed an increased mitotic index for gene knockouts targeting established components of the kinetochore and mitotic spindle. In contrast, we observed a reduced mitotic index for components of the spindle assembly checkpoint, including MAD2L1, BUB1B, and TTK (Mps1).

**Fig. 4.**
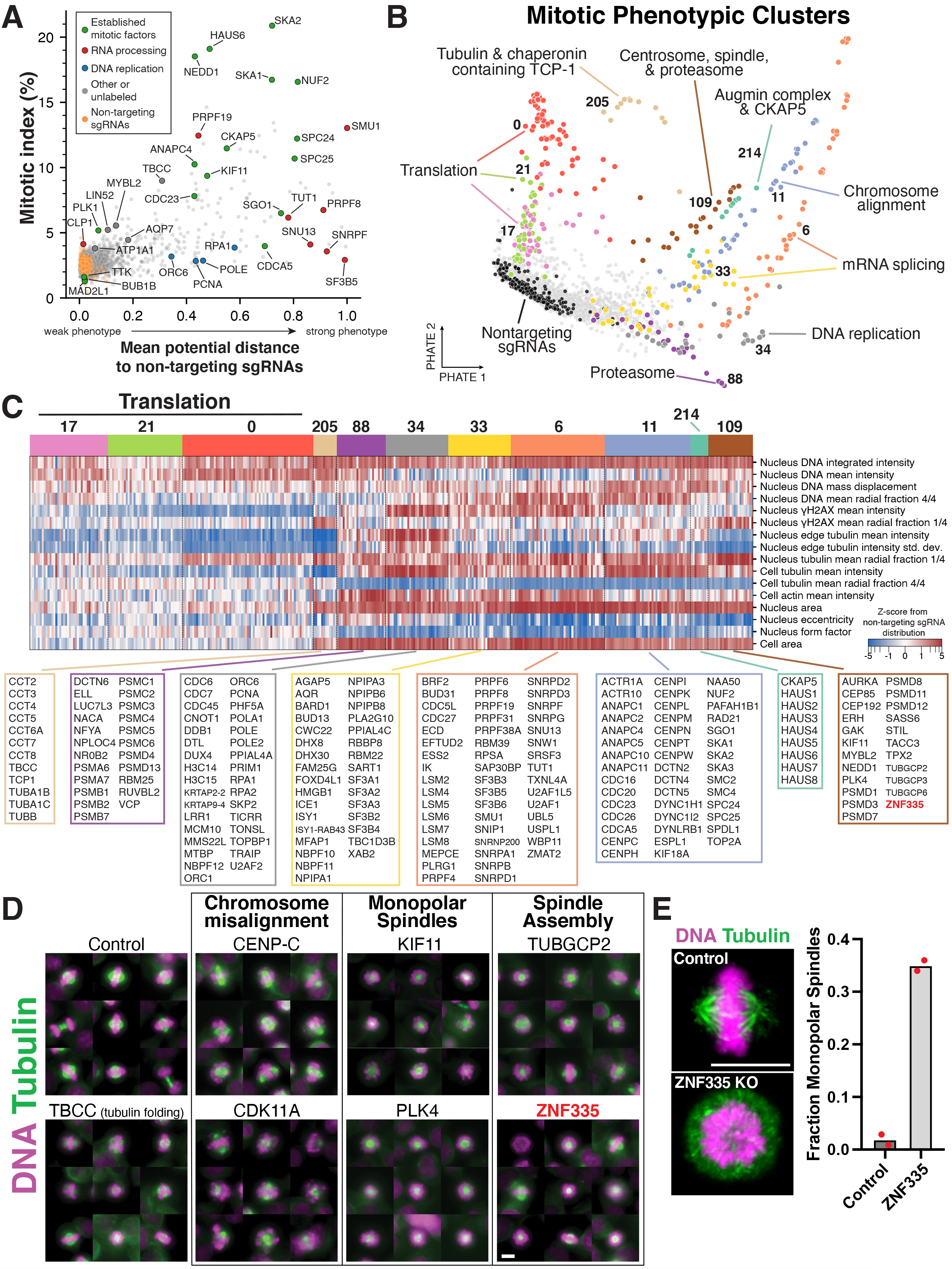
Mitotic phenotypes uncover essential genes required for cell division. (**A**) Scatter plot showing mitotic index (proportion of mitotic cells) of the imaged cell population for each gene target compared to a summary score of mitotic phenotype strength computed by PHATE (*35*) high-dimensional analysis. The summary phenotype score is the mean PHATE potential distance to non-targeting control sgRNAs for each gene, normalized between 0 and 1. This highlights the complementary but distinct information provided by computational analysis of image-based mitotic phenotypes. Labeled gene targets are colored by functional category and highlight factors with increased cell division defects. (**B**) Two-dimensional representation of the mitotic phenotype landscape of gene targets in the primary screen computed using PHATE with hundreds of summary phenotype parameters, and then clustered to form groups of genes with similar phenotypes (see Methods). Each dot represents a single gene target, colored corresponding to the indicated cluster. Functional descriptions of clusters summarize the roles of the contained gene targets. (**C**) Heat map of mitotic knockout phenotypes corresponding to the clusters in (B) for a manually-selected subset of phenotype parameters. This highlights the phenotypic similarity of gene targets within clusters and the ability to separate distinct mitotic functions. Furthermore, this analysis implicated the potential role of ZNF335 in mitotic spindle function. All genes from selected clusters are listed below. Parameters are presented as z-scores from the distribution of non-targeting sgRNAs, visualized on a symmetric log scale (linear between -1 and 1). (**D**) Example images of mitotic cells from the screen visualizing DNA (magenta) and microtubules (green) from gene targets selected in the computational and/or visual analysis highlight the diversity of identified mitotic phenotypes (see also Fig. S7C). Scale bar, 10 µm. (**E**) Left, immunofluorescence images showing individual cell lines stably expressing a single control sgRNA or a sgRNA targeting ZNF335. Right, bar plot of the corresponding fraction of mitotic cells with monopolar spindles; each data point represents one experiment with >100 cells. This demonstrates the reproducible strong effect of knocking out ZNF335 on spindle assembly. Images are deconvolved maximum intensity projections of fixed cells stained for microtubules (anti-alpha-tubulin) and DNA (Hoechst). Scale bars, 10 µm.

Similar to our analysis of interphase cells, we next selected 876 non-redundant measurements from the extracted image parameters of mitotic knockout cells, including the overlap of tubulin and DNA staining as a measure of mitotic chromosome alignment, and clustered gene targets with similar phenotype profiles (Fig. 4B; Fig. S4D-F; Methods). In addition, we conducted a manual visual analysis for each gene, with two individuals blindly and independently scoring image montages for the presence of mitotic defects. Overall, we found a strong correspondence between the automated and manual scoring (Fig. S7A, B). This combined analysis identified a wide range of established players in cell division, including factors with roles in cell cycle control, mitotic spindle assembly, kinetochore function, sister chromatid cohesion, and cytokinesis (Fig. 4B-D; Fig. S7C). From our computational analysis, we identified multiple mitotic clusters with functionally-related genes (Fig. 4B, C), despite decreased cell counts and increased morphological heterogeneity as compared to interphase cells. For example, we observed close clustering of CKAP5 and the entire Augmin complex (mitotic cluster M214), tubulin subunits with tubulin folding factors and the CCT chaperonin (M205), DNA replication factors (M34), factors required for chromosome alignment including kinetochore components (M11), and spindle and centrosome components (M109). In addition, we identified clusters for gene targets with established roles in mRNA splicing (M6 and M33), proteasome function (M88), and ribosome function (M0, M17, and M21) indicating that mitotic phenotype parameters are able to distinguish these functional categories. We found that this high-dimensional computational analysis provided a complementary but distinct measurement of mitotic phenotypes as compared to mitotic index (Fig. 4A). Visual analysis of cell image montages further allowed us to distinguish individual gene targets for their specific roles during mitosis (Fig. 4D; Fig. S7C). For example, we were able to detect reduced microtubule density following depletion of the tubulin chaperone TBCC, chromosome mis-alignment following depletion of kinetochore components (e.g., CENP-C and Ska1) and additional factors (SMU1 and CDK11A, amongst many others), monopolar spindles associated with the depletion of Kif11 or Plk4, and short mitotic spindles in knockouts of CKAP5 or Augmin subunits (e.g., HAUS6).

Predicted roles in mitosis also emerged for poorly characterized genes based on co-clustering of genes with well-defined mitotic functions. In particular, we found that ZNF335 clustered with spindle proteins (including TACC3 and TPX2), gamma-tubulin complex proteins (TUBGCP2/3/6), and some proteasome (PSMD) subunits (cluster M109; Fig. 4C). Examination of mitotic cells lacking ZNF335 suggested the presence of spindle defects (Fig. 4D). We confirmed this phenotype by generating a stable inducible knockout cell line for ZNF335, which displayed a substantially increased proportion of cells with monopolar spindles (Fig. 4E). Together, this suggests a role for ZNF335 in centrosome and spindle function, providing a potential explanation for why ZNF335 mutations are observed in human microcephaly (*47*), as is the case for many centrosome components (*48*). In addition, we identified dozens of genes with mitotic phenotypes, but less clear phenotypic clustering, that have not been implicated previously as having roles in cell division (see below). Together, this analysis validates pooled large-scale image-based screening to identify complex interphase and mitotic phenotypes.

## A pooled live-cell imaging-based screen for mitotic defects

Based on the large number of genes with unexpected mitotic phenotypes and the power of microscopy-based approaches to directly visualize these phenotypes in detail (*49, 50*), we performed a secondary pooled live-cell screen to analyze these factors further. First, we defined a list of 229 genes with unexpected mitotic phenotypes, and additionally selected 10 positive control genes with established roles in diverse mitotic processes. We generated a lentiviral library of Cas9 guides targeting these genes, with 2 sgRNAs per gene and 50 non-targeting sgRNAs (526 total sgRNAs; see Methods). The sgRNA library was transduced into a HeLa cell line containing doxycycline-inducible Cas9 and a constitutively-expressed H2B-mCherry fusion to visualize chromatin (Fig. 5A). We conducted time-lapse imaging of the pooled cell population for 24 hours with time points at 10 minute intervals, after either 48 or 72 hours of Cas9 expression in separate experiments. Following the acquisition of the time-lapse images, we immediately fixed the cell population and amplified the sgRNA sequences *in situ* to identify the gene targeted in each cell, as described for the fixed-cell screen (Fig. 5A; Methods). After tracking cell lineages through each time course and using a support vector classifier to identify mitotic cells, we obtained time-lapse movies for 451,434 total cell division events, with a median of 1,381 division events per gene target (Fig. S8A-C; Data S3; Methods). This enabled us to generate time-lapse montages of tracked cells for each gene target, with each cell temporally aligned to mitotic entry (Fig. 5B-C; Movie S1; https://nematode.wi.mit.edu/vesuvius/). Using this approach, we observed the expected phenotypes for positive controls (Fig. 5B), indicating that optical pooled screening provides an effective and scalable strategy for live-cell screening of complex cell biological phenotypes.

**Fig. 5.**
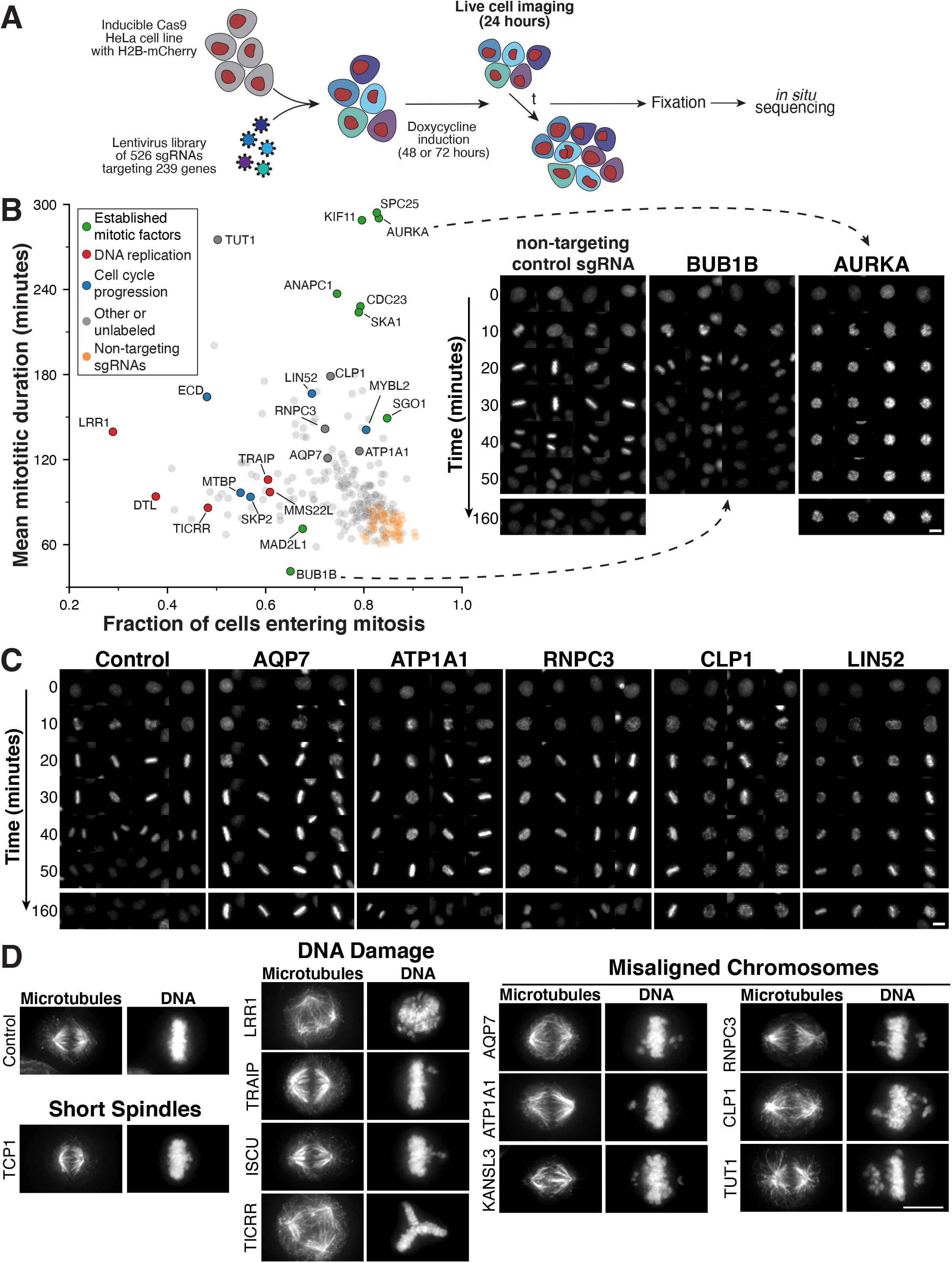
A pooled live-cell screen identifies gene targets required for mitotic progression. (**A**) Schematic of the experimental workflow for the live-cell, image-based pooled CRISPR screen using a cell line expressing an H2B-mCherry fusion (also see Methods). (**B**) Left, scatter plot comparing the fraction of cells that enter mitosis within the 24 hour time course and the mitotic duration of observed cell division events. This live-cell analysis identifies genes with defects in mitotic entry or progression. Labeled genes are colored by functional category. Right, example images of H2B-mCherry fluorescence from the live-cell screen at the indicated time points after mitotic entry for knockouts of established cell division components. Targeting the spindle assembly checkpoint gene BUB1B results in an acceleration of mitosis, whereas knockouts of AURKA, a key mitotic kinase, result in a mitotic arrest. (**C**) Example time course montages from the live-cell screen as in (B) demonstrating mitotic delay and mitotic defects for selected target genes. (**D**) Immunofluorescence images showing individual cell lines stably expressing a single sgRNA targeting each gene of interest to enable visualization of phenotypes at higher resolution across a single population (see also Fig. S9A). Images are deconvolved maximum intensity projections of fixed cells stained for microtubules (anti-alpha-tubulin) and DNA (Hoechst). Scale bars, 10 µm.

Using this automated live-cell analysis of cell division, for each gene we calculated the mean duration of division events, as well as the fraction of cells that enter mitosis during the time course (Fig. 5B). As expected based on the presence of mitotic defects in the primary screen, the majority of gene knockouts displayed an increased mitotic duration relative to non-targeting controls. In contrast, we detected a decreased mitotic duration for the established spindle assembly checkpoint component BUB1B. We were also able to distinguish gene targets with established or predicted roles in DNA replication or repair (e.g., DTL, LRR1, TICRR, MMS22L), based on their increased mitotic duration but reduced fraction of cells entering mitosis (Fig. 5B), indicative of defective mitotic entry. We next visually inspected each time-lapse montage for the presence of mitotic phenotypes, including lagging chromosomes or delayed chromosome alignment (Fig. 5C; Fig. S8D; Data S3). From these observed phenotypes, we selected 29 genes of interest to conduct targeted downstream analyses. In each case, we generated individual cell lines with a single sgRNA targeting the corresponding gene and conducted both fixed- and live-cell microscopy to identify phenotypes for each individual gene knockout at higher spatial and temporal resolution (Fig. 5D; Fig. S9A; Movie S2). The majority of the selected gene targets displayed clear defects in chromosome alignment and segregation. Of these genes, a subset displayed defects in bipolar spindle assembly, including short spindles (TCP1). We were also able to distinguish gene targets with roles in DNA replication or DNA damage (e.g., LRR1, TRAIP, ISCU, TICRR, MMS22L) which resulted in multipolar spindles or misaligned chromosomes (Fig. 5D; Fig. S9A). Unexpectedly, amongst the gene targets whose knockouts resulted in misaligned chromosomes, we identified two membrane-bound transporters - the plasma membrane-localized aquaporin AQP7 and the sodium/potassium-transporting ATPase ATP1A1 (Fig. 5D). AQP7 is the only aquaporin that is broadly essential for cellular viability in the DepMap database (*12*). AQP7 and ATP1A1 knockouts both displayed a reproducible delay in chromosome alignment and an extended mitotic duration, but we did not observe defects in bipolar spindle assembly (Fig. 5D) or kinetochore function (Fig. 6A, B). We propose that AQP7 and ATP1A1 are required to create an internal cellular environment that promotes proper chromosome segregation.

**Fig. 6.**
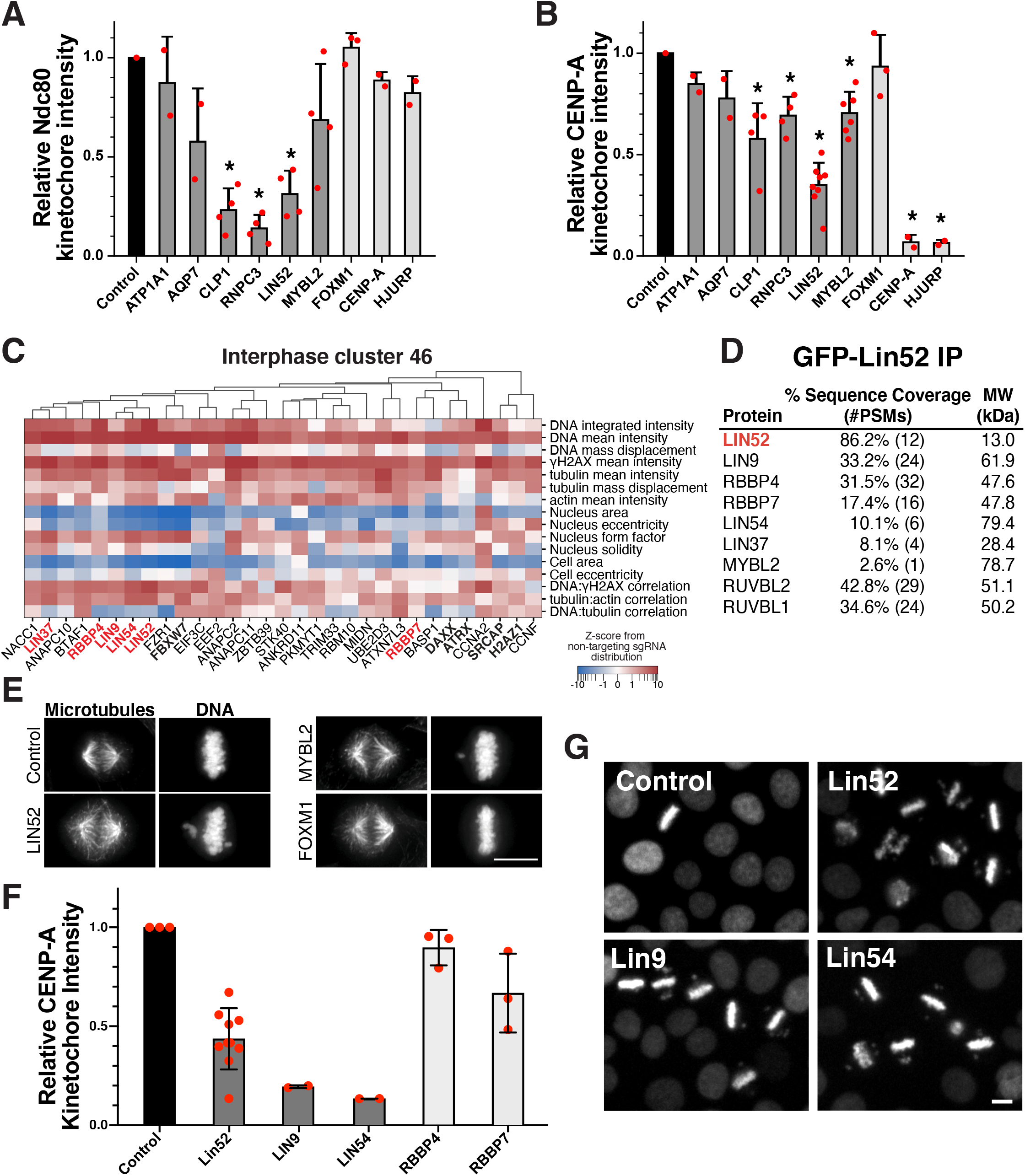
The Lin52 complex functions to promote proper kinetochore assembly and chromosome segregation. (**A**) Bar plot showing kinetochore-localized intensity of the outer kinetochore microtubule-binding protein Ndc80 in the indicated inducible knockout cell lines. Ndc80 levels are substantially decreased for CLP1, RNPC3, and LIN52 knockouts, but not for other hit genes from the live-cell pooled screen. Each data point represents the median kinetochore signal of one experiment for >10 cells per gene target. Values are normalized relative to control cells from the same experiment. *P<0.01 by two-tailed independent T-test relative to control cells. (**B**) Bar plot showing kinetochore-localized intensity for the inner kinetochore centromere-specific histone CENP-A in the indicated inducible knockout cell lines. CENP-A intensity is significantly decreased for LIN52. Experiment design as in (A). *P<0.01 by two-sided independent T-test relative to control cells. (**C**) Heat map of primary screen interphase cluster 46 knockout phenotypes for a manually-selected subset of parameters. This highlights the similarities between LIN52 and its previously defined interacting partners (red) with potential co-functional chromatin factors (bold). Hierarchical clustering (top) was performed using the correlation of high-dimensional phenotype profiles (see Methods). Parameters are presented as z-scores from the distribution of non-targeting sgRNAs, visualized on a symmetric log scale (linear between -1 and 1). (**D**) Mass spectrometry analysis of an immunoprecipitation of GFP-LIN52 from mitotically-enriched human cells relative to controls, indicating that LIN52 associates with a subset of expected factors, but not the entire DREAM complex. (**E**) Immunofluorescence images of microtubules (anti-alpha-tubulin) and DNA (Hoechst) in inducible knockout cell lines identify chromosome alignment defects for RUVBL1, RUVBL2, and the DREAM complex components LIN52 and MYBL2, but not FOXM1. (**F**) Bar plot showing kinetochore-localized intensity for the inner kinetochore centromere-specific histone CENP-A as in (B). LIN52, LIN9, and LIN54 each demonstrate a substantial decrease in CENP-A kinetochore localization. (**G**) Images from time-lapse fluorescence imaging of individual knockout cell lines expressing H2B-mCherry, demonstrating similar strong mitotic phenotypes for LIN52, LIN9, and LIN54. Also see Movie S3. Scale bars, 10 µm.

To define the basis for observed mitotic phenotypes, we next tested the function of the kinetochore, the key player in mediating interactions between centromere DNA and microtubule polymers during cell division (*51*). In particular, we tested the localization of the inner kinetochore centromere-specific histone CENP-A and the outer kinetochore microtubule-binding protein Ndc80. Of the 29 gene targets tested, the majority displayed only modest changes in the kinetochore recruitment of Ndc80, suggesting that kinetochore assembly is largely normal. In contrast, we observed a substantial reduction in Ndc80 localization for the CLP1, RNPC3, and LIN52 inducible knockouts (Fig. 6A; Fig. S9B). Similarly, we found that most knockouts did not strongly alter the levels of CENP-A, with the exception of the LIN52 inducible knockout (Fig. 6B; Fig. S9C). Importantly, established knockouts that prevent proper CENP-A recruitment, including HJURP and CENP-A itself (*52*), do not affect Ndc80 recruitment at the tested time points (Fig. 6A, B), suggesting that LIN52 is required for a process that contributes to both inner and outer kinetochore assembly. Thus, CLP1 and RNPC3 are required for robust outer kinetochore recruitment, likely through their established roles in mRNA processing, whereas LIN52 depletion compromises multiple aspects of kinetochore assembly.

To determine the basis for the phenotypes observed in LIN52 knockouts, we next considered its functional relationships. Prior work found that LIN52 is a component of the DREAM complex, comprised of E2F family transcription factors, LIN9/37/52/54, MYBL1/2, RBL1/2, RBBP4, and TFDP1/2, which acts together with FOXM1 as a transcriptional regulator for multiple cell cycle genes (*53*). In contrast, in our analysis of interphase phenotypes from the primary screen, LIN52 clustered with LIN9/37/54, RBBP4 and RBBP7 (cluster 46; Fig. 6C), but not with the other DREAM-related genes analyzed in our fixed-cell screen (MYBL2, TFDP1, E2F1, E2F3, E2F6, FOXM1). Similarly, in immunoprecipitation-mass spectrometry experiments we found that LIN52 associated with LIN9/37/54, RBBP4, and RBBP7, but not other established DREAM complex proteins (Fig. 6D). Prior work has implicated RBBP4 and RBBP7 (also known as RbAp46 and RbAp48) in promoting centromere chromatin formation (*54*), but the basis for this role is unclear. Consistent with the phenotypic co-clustering and physical interactions, we observed chromosome misalignment, a mitotic delay, and substantial changes to kinetochore assembly in knockouts of either LIN52, LIN9, or LIN54 (Fig. 6A, B, E, F, G; Fig. S9B, C; Movie S3). In contrast, we did not detect altered kinetochore protein levels or chromosome misalignment for FOXM1 knockouts and only observed a modest change in CENP-A and Ndc80 localization in MYBL2 knockouts (Fig. 6A, B, E). RNA-sequencing analysis of LIN52, FOXM1, and MYBL2 knockout cell lines identified partially-overlapping sets of differentially-expressed genes compared to a control sgRNA, but did not identify pervasive changes in the expression of centromere components that would explain the observed phenotypes (Fig. S9E). Thus, the combination of the physical and functional interactions that we observed for LIN52 define a new functional module comprised of at least LIN52, LIN9, LIN37, and LIN54 that plays a role in centromere function, possibly through changes to underlying chromatin that promote the localization of both inner and outer kinetochore components.

## Pooled image-based screens define the phenotypic landscape of cellular functions

Together, our pooled microscopy-based analysis of tens of millions of individual knockout cells for thousands of essential and fitness-conferring human genes defines their functional contributions to diverse biological processes. By obtaining quantitative information for diverse image-based parameters that are directly comparable across a large cell population, this approach identifies co-functional gene targets with fine-grained resolution to distinguish roles in specific cellular processes and protein complexes. Studies analyzing proteome-wide protein interactions and coordinated gene expression across biological contexts have previously defined large-scale molecular networks. The precision and breadth of the clustering behaviors reported here highlight the ability of quantitative image-based phenotypic profiling to provide a similar scale of functional information, with complementary but distinct insights. Although we focused primarily on aggregate behaviors across the population of imaged cells for each gene target, future studies leveraging the distribution of single-cell phenotypes will enable additional resolution and insights for understanding gene functions. In addition to providing an expansive and powerful resource for the analysis of phenotypes resulting from the disruption of essential genes in our companion interactive web portal (https://nematode.wi.mit.edu/vesuvius/), this work provides multiple predictions for the contributions of incompletely characterized genes to fundamental cellular functions. We anticipate that this type of scalable and cost-effective cell biological genomic screening will enable future studies that will yield additional key insights across numerous cellular phenotypes and conditions.

## ACKNOWLEDGEMENTS

We thank Andy Nutter-Upham and Scott McCallum for creating the Vesuvius web portal, members of the Blainey and Cheeseman labs, Bingbing Yuan and Heather Keys for their support and input, and Dave Bartel, Kara McKinley, Anne Carpenter, Anna Le, and Russell Walton for comments on the manuscript. We thank the Broad Genomic Perturbation Platform for providing CFD scores for our custom sgRNA library and Celeste Diaz and Julia Bauman in the lab of J.T. Neal at the Broad Institute for assistance in developing custom antibody conjugations. The HeLa cell line was used in this research. Henrietta Lacks, and the HeLa cell line that was established from her tumor cells without her knowledge or consent in 1951, have made significant contributions to scientific progress and advances in human health. We are grateful to Lacks, now deceased, and to the Lacks family for their contributions to biomedical research.

## Author contributions

Conceptualization: LF, KCS, DF, AS, PCB,IMC;Methodology,Validation,Investigation:LFforscreen experiments, KCS for all other experiments; Formal Analysis: LF for computational screen analysis and RNA-sequencing data, KCS for all other data, KCS & BM for visual annotation of screen images; Writing - Original Draft Preparation: LF, ICM; Writing - Review & Editing: LF, KCS, DF, AS, PCB, IMC; Supervision: PCB, IMC; Funding Acquisition: IMC, PCB.

## Data and materials availability

Processed images and data from the screen are available in the supplemental materials and through the companion web portal (https://nematode.wi.mit.edu/vesuvius/), with raw image data available upon request. Public code repositories are available containing functions used for processing optical pooled screening data (https://github.com/feldman4/OpticalPooledScreens) and application-specific tools (https://github.com/lukebfunk/OpticalPooledScreens). RNA-seq data are deposited in the Gene Expression Omnibus: GSE188631.

## MATERIALS AND METHODS

### Library design and cloning

The primary screen library of fitness-conferring genes was defined based on evidence from multiple published sources. First, we used data from the Broad Institute DepMap project (*11, 12, 15*) to identify genes that are broadly fitness-conferring in a variety of cell lines. Specifically, we selected genes with a genetic dependency probability of >0.35 in at least 10% of the >600 tested cell lines (Fig. S1A, B), resulting in 3,991 selected genes. We subsequently chose 1,081 additional genes that had evidence of essentiality in at least 2 other published screens (*10, 13, 14, 16*–*18*). CRISPR sgRNA sequences were selected from published libraries (*18*–*20*), with simultaneous optimization of sgRNA performance (e.g., on- and off-target efficiency) and minimization of 5’ sequence length required to demultiplex all sgRNAs during in situ sequencing. In total, we selected 20,445 sgRNA sequences, including 4 sgRNAs each for the vast majority of gene targets and 250 non-targeting control sgRNAs, with a minimum Levenshtein distance of 2 between the leading 11-nucleotide 5’ sequence for all possible pairs of sgRNAs (Data S1). We note that, for some groups of genes with high sequence homology, it is not possible to design distinct targeting sgRNAs for each gene. For groups of genes where the full lists of possible sgRNAs collected from previously published libraries were identical, a single set of 4 sgRNAs was chosen to target these genes collectively. Two sgRNAs per gene were selected for the 239 genes in the live cell screen based on performance in the fixed-cell screen, in addition to 50 non-targeting guides selected using the 5’ sequence optimization described above. Targeting and non-targeting sgRNA libraries were designed as separate subpools of synthesized oligo arrays (Agilent) and independently cloned into CROPseq-puro-v2 (Addgene #127458) as described previously (*9*).

For expression of fusion proteins, H2B (pKC96) was amplified from a template retroviral construct (*55*), while C7orf26 (NP_076972.2; pKC509) and LIN52 (Q52LA3.1; pKC518) were human codon-optimized and synthesized (Twist Biosciences). Gene fragments were ligated into either an mCherry or EGFP pBABE-based vector (Addgene #44432). sgRNA constructs for individual inducible knockout cell lines were generated by primer annealing and ligation into sgOPTI (*56*; see Data S4). A control sgRNA with a single target site within the non-essential LBR gene was used for comparison of all follow-up experiments (HS1, *57*).

### Tissue culture

For a list of cells used in this study, see Data S4. HeLa and HEK293 cells were cultured in DMEM with sodium pyruvate and GlutaMAX (Life Technologies 10569044) or 2 mM L-glutamine supplemented with 10% heat-inactivated fetal bovine serum (Sigma F4135) and 100 U/mL penicillin-streptomycin (Thermo Fisher Scientific 15140122).

### Virus production and transduction

Prior to lentiviral production of screening sgRNA libraries, the corresponding targeting and non-targeting plasmid pools were mixed (final non-targeting sgRNA pool fraction of 5% for the primary fixed-cell screen, 9.5% for the secondary live-cell screen). Lentiviral production and transduction were performed as described previously for libraries (*8, 9*) or single targets (*17*). Retrovirus was generated by transfecting VSVG packaging plasmid and pBABE-based vectors containing H2B-mCherry, EGFP-C7orf26, or EGFP-Lin52 fusions into HEK293-GP cells with Effectene (Qiagen) for transduction as described previously (*58*).

### Fluorescence microscopy

All screening datasets were acquired using a Nikon Ti-2 inverted epifluorescence microscope with automated stage control, hardware autofocus, and an Iris 9 sCMOS camera (Teledyne Photometrics). All hardware was controlled using NIS-Elements AR, and a CELESTA light engine (Lumencor) was used for fluorescence illumination. *In situ* sequencing cycles were acquired using a 10X 0.45 NA CFI Plan Apo Lambda objective (Nikon MRD00105) and 2×2 pixel binning with the following laser lines, filters, and exposure times for each channel: DAPI (408 nm laser excitation with 0.8% power, custom Chroma dual-band 408/473 dichroic and emission filter set, 50 ms exposure), Miseq G (545 nm laser with 30% power and Semrock FF01-543/3 excitation filter, Chroma T555LPXR dichroic filter, Chroma ET575/30 emission filter, 200 ms exposure), Miseq T (545 nm laser excitation with 30% power, Chroma T565LPXR dichroic filter, Semrock FF01-615/24 emission filter, 200 ms exposure), Miseq A (635 nm laser excitation with 30% power, Chroma ZET635RDC dichroic filter, Semrock FF01-680/42 emission filter, 200 ms), Miseq C (635 nm laser excitation with 30% power, Chroma ZET635RDC dichroic filter, Semrock FF01-732/68 emission filter, 200 ms exposure). Fixed-cell primary screen phenotype images were acquired using a 20X 0.75 NA CFI Plan Apo Lambda objective (Nikon MRD00205) using DAPI (as before), FITC (473 nm laser excitation, custom Chroma 408/473 filter set), Alexa Fluor 594 (same settings as MiSeq T), and Alexa Fluor 750 (750 nm laser excitation, Semrock FF765-Di01 dichroic filter, custom ET820/110 Chroma emission filter) fluorescence channels. For the live-cell secondary screen, timelapse phenotype images were acquired using the 20X objective lens, an mCherry fluorescence channel (same settings as MiSeq T), and a microscope enclosure with temperature and CO_2_ control along with passive humidification (Okolab H201).

Immunofluorescence images of single knockout cell lines were taken on the Deltavision Ultra (Cytiva) system using a 60x/1.42NA objective and deconvolution. For kinetochore component quantification, z-sections at 0.2 μm intervals were taken using a 100X/1.45NA objective. For time lapse imaging of individual inducible knockouts and EGFP fusion cell lines, we used a Nikon Eclipse microscope equipped with an ORCA-Fusion BT sCMOS camera (Hamamatsu) using a Plan Fluor 20X/0.5 NA (live cells) or 40x/1.3NA (EGFP) objective lens.

### Fixed-cell optical pooled CRISPR screen

For the fixed cell screen, HeLa-TetR-Cas9 (A7) cells were transduced with the 20,445 sgRNA library in CROPseq-puro-v2 and selected with 2 μg/mL puromycin (Thermo Fisher Scientific A1113803) for 4 days. Cas9 expression was induced with 2 μg/mL doxycycline for 78 hours, and the cell library was seeded into eight 6-well glass-bottom plates (Cellvis P06-1.5H-N) at a density of 300,000 cells per well (∼30,000 cells/cm^2^) 48 hours prior to fixation. Cells were fixed with 4% paraformaldehyde in PBS for 30 minutes, followed by *in situ* amplification as described previously (*8, 9*). After rolling circle amplification, cells were stained with rabbit anti-gamma H2A.X (phospho S139) antibody (Abcam ab81299, 1:2000 dilution in PBS with 3% BSA) for 1 hour at room temperature. Cells were washed twice with PBS-T (PBS with 0.05% Tween-20), then stained with mouse anti-alpha-tubulin-FITC antibody (Sigma F2168, 1:500 dilution), goat anti-rabbit antibody disulfide-linked to Alexa Fluor 594 (Invitrogen 31212, Thermo Fisher Scientific A10270, custom conjugation; 1:500 dilution), and Alexa Fluor Plus 750 Phalloidin (Thermo Fisher Scientific A30105, 1:1000 dilution) in PBS with 3% BSA for 45 minutes at room temperature. After washing with PBS-T three times, well plates were replaced with 200 ng/mL DAPI in 2X SSC and imaged for cellular phenotypes using the microscope configuration described above with 4 z-slices at 1.5 μm intervals. Following phenotype imaging, Alexa Fluor 594 was cleaved from disulfide-linked antibodies by incubating cells in 50 mM TCEP in 2X SSC for 1 hour at room temperature, followed by three washes with PBS-T. Finally, 11 cycles of *in situ* sequencing-by-synthesis were performed as described previously (*8, 9*). In parallel with the optical pooled screen, cells expressing the same sgRNA library were induced with 1 μg/ mL doxycycline, and then doxycycline media was refreshed every day for 2 more days. Cells were harvested on days 0 (pre-induction), 3, and 5 post-Cas9 induction and genomic DNA was extracted using PureLink (Invitrogen). sgRNA sequences were then PCR amplified using Q5 hotstart (NEB) with primers oDF344 (ACACGACGCTCTTCCGATCTtcttgtggaaaggacgaaac) and oDF112 (CTGGAGTTCAGACGTGTGCTCTTCCGATCaagcaccgactcggtgccac) before addition of index barcodes and sequencing on an Illumina HiSeq using sequencing primer oKC651 (ACACTCTTTCCCTACACGACGCTCTTCCGATCTtcttgtggaaaggacgaaacaccg).

### Live-cell optical pooled CRISPR screen

HeLa-TetR-Cas9 cells expressing an H2B-mCherry fusion protein (cKC556) were transduced with the live-cell screening library of 526 sgRNA sequences. Cells were selected with 2 μg/mL puromycin (Thermo Fisher Scientific A1113803) for 3 days. Cas9 expression was induced with 2 μg/mL doxycycline for 48 (day 3 time course) or 72 hours (day 4 time course) prior to the beginning of live-cell imaging, and cells were seeded into 6-well glass-bottom plates (Cellvis P06-1.5H-N) at a density of 300,000 or 350,000 cells per well (∼35,000 cells/cm^2^) 24 hours prior to imaging. Each time course was performed in three batches on separate days. Immediately before imaging, cells were washed once with PBS, and then replaced with imaging media consisting of phenol red-free DMEM with L-glutamine and HEPES (Thermo Fisher Scientific 21063029) supplemented with 10% heat-inactivated fetal bovine serum (Sigma F4135) and 100 U/mL penicillin-streptomycin (Thermo Fisher Scientific 15140122). Live-cell imaging was performed using the microscope configuration described above and 2 z-slices spaced at either 4 or 5 μm intervals. Cells were imaged for 24 hours at 10 minute time intervals, immediately fixed with 4% paraformaldehyde in PBS for 30 minutes, then processed through *in situ* amplification and sequencing-by-synthesis following the same protocol as the fixed-cell screen.

### Screening image analysis

*In situ* sequencing spots were identified and barcode sequences extracted using our previously described workflow (*8, 9, 59*). In addition, phenotype images were acquired at a higher magnification than *in situ* sequencing images, and thus the datasets were computationally aligned to match cell identities. This alignment was completed by computing the Delaunay triangulation of nuclei centroids for each phenotype and sequencing image tile, and then comparing triangulations between images from the two datasets to find matching tiles and cell identities.

Phenotype images from both screens were first maximum intensity projected to compress z-slices into a single plane, and then a retrospective flat-field correction was applied to reduce effects from uneven illumination (*60*). Nuclei were semantically segmented by applying a local intensity threshold to the DAPI channel and then performing morphological operations to remove aberrant holes and particles. Individual nucleus instances were then segmented using the watershed algorithm. In the fixed-cell phenotype data, semantic segmentation of cytoplasmic foreground was achieved by thresholding a Gaussian-filtered copy of the phalloidin (actin) channel (sigma of 3 pixels), followed by morphological operations. Cell instances were identified by applying the watershed algorithm with nuclear segmentations as seeds. Phenotype parameters were extracted from nuclear and cellular segmentations for each channel by implementing CellProfiler-based features and additional measurements as Python functions operating on scikit-image RegionProperties objects (*61*). Image segmentation, phenotype feature extraction, and *in situ* sequencing analysis were performed in parallel on a per-image tile basis using the Snakemake workflow manager (*62*).

### Fixed-cell screen phenotype analysis

After aligning the phenotype and sequencing datasets, a subset of features were transformed to approximate normal distributions (see Data S2). All features for each cell were then normalized using the median and median absolute deviation of the population of cells carrying non-targeting sgRNAs within the same well (robust z-score). This internal control procedure was used to reduce batch effects between wells and plates that may be caused by intensity differences or cell density effects. Mitotic and interphase cells were identified using a support vector classifier (scikit-learn (*63*) svm.SVC implementation, default parameters) with a subset of 182 features (Fig. S1F; Data S2). Cell-level measurements were then re-normalized from the raw data as before, but within interphase and mitotic cells separately.

Summary phenotype measurements were computed for each gene target by taking the median of z-scored parameters for all cells targeted by a single sgRNA sequence, then aggregating to the gene level by taking the median across sgRNAs targeting the same gene. Raw p-values for a subset of summary parameters were computed by comparing gene scores to null distributions of corresponding bootstrapped summary scores from cells expressing non-targeting sgRNAs. Separate null distributions were defined for each gene target by first performing 100,000 cell sampling repetitions to produce a distribution of bootstrapped non-targeting sgRNA scores for each cell sample size of the targeting sgRNAs. These guide-level null distributions were then correspondingly sampled 100,000 times for each group of sgRNAs targeting the same gene and aggregated to produce gene-level null distributions with matched cell and guide sample sizes. The Benjamini-Hochberg procedure was applied to obtain the reported FDR q-values. An FDR threshold of 0.05 was used for defining significance for all parameters, with effect size thresholds additionally used for mean nuclear γH2AX intensity, set by the 2.5 and 97.5 percentiles of the non-targeting sgRNA scores.

For the high-dimensional analysis, pairs of features with a Pearson correlation greater than 0.9 were iteratively excluded and additional features known to be uninformative were removed, resulting in a set of 472 features for the interphase dataset and 884 features for the mitotic dataset. Further feature redundancies were reduced by applying principal component analysis (PCA) and retaining the components that explain 95% of the variance in the datasets (103 components for the interphase data, 530 components for the mitotic data). The PHATE manifold learning and visualization algorithm (*35*) was then used to produce two-dimensional representations of the phenotypic landscape of gene targets (default parameters except n_pca=None). To cluster knockout phenotypes, the PHATE diffusion operator affinity graph was supplied as input to the Leiden algorithm, which optimizes cluster modularity (*64*). The Leiden resolution parameter was chosen by analyzing the robustness of clustering solutions to the subsampling of gene-level data with varying resolution (resolution = 10 for interphase dataset, resolution = 9 for mitotic dataset; Fig. S4C, F). In parallel with the computational phenotype analysis, two individuals independently scored mitotic phenotypes from the primary screen by visually inspecting montages of mitotic cells from each gene target and assigning a phenotype severity score from 1 to 9 (Fig. S7A, B). During this process, the scoring individuals were blinded to the gene identities associated with each montage of cells.

### Live-cell screen phenotype analysis

Following nuclear segmentation of the time lapse data, cells were tracked across frames using the TrackMate implementation of the linear assignment problem approach to particle tracking (*65, 66*). The cost of linking nuclei in consecutive frames was set as the squared distance between centroids, with maximum linking distance set to 60 pixels (∼18 μm). Track gaps up to 2 frames were allowed, in addition to track merges and splits. Tracked cell lineages that did not last for the full 24 hour time-course were excluded from analysis.

The sgRNA assigned to each tracked cell lineage in the phenotype data was determined by matching cell identities between the *in situ* sequencing images and the final time point of the time course. Similar to the fixed-cell screen, individual cell feature measurements were normalized using the median and median absolute deviation of the non-targeting control cell population from the same well and time point to reduce batch effects and correct for temporal intensity variations. Interphase, mitotic, and apoptotic cells were classified using a support vector machine (scikit-learn svm.SVC, linear kernel) with a subset of 81 features (Fig. S8A; Data S3). However, due to the difficulty of separating mitotic and apoptotic cells based on H2B-mCherry fluorescence alone, these categories were later combined into a single, broad “mitotic” bin. Cell division events were defined as a contiguous sequence of at least 2 frames of mitotic-classified cells immediately followed by a split in the track into 2 daughter cells (Fig. S8B). Also included as cell division events were continuous sequences of mitotic cells that start in the first frame or reach the end of the acquired time course, if the observed mitotic duration was at least as long as the average mitotic duration of non-targeting control cells in the same well. Mitotic duration was measured as the time difference between the first and last frame of the cell division event. The fraction of cells entering mitosis was calculated as the fraction of tracked lineages containing at least one cell division event as defined above. Since many genes exhibited a stronger phenotype at either the Day 3 or Day 4 time point, likely due to differences in protein depletion timing, the strongest phenotype was selected for plotting in Fig. 5B by selecting the time course with the highest absolute difference in mitotic duration compared to the mean of non-targeting sgRNAs.

### GFP immunoprecipitation and Mass-spectrometry

IP-MS experiments were performed as described previously (*67*). EGFP-C7orf26 and EGFP-LIN52 cells were mitotically enriched with 10μM STLC overnight, harvested and washed in PBS and resuspended 1:1 in 1X Lysis Buffer (50 mM HEPES, 1 mM EGTA, 1 mM MgCl_2_, 100 mM KCl, 10% glycerol, pH 7.4) then frozen in liquid nitrogen. Cells were thawed after addition of an equal volume of 1.5X lysis buffer supplemented with 0.075% Nonidet P-40, 1X Complete EDTA-free protease inhibitor cocktail (Roche), 1 mM phenylmethylsulfonyl fluoride, 20 mM beta-glycerophosphate, 1 mM sodium fluoride, and 0.4 mM sodium orthovanadate. Cells were then lysed by sonication and cleared by centrifugation. The supernatant was mixed with Protein A beads (Biorad) coupled to rabbit anti-GFP antibodies (Cheeseman lab) and rotated at 4°C for 1 hour. Beads were washed five times in wash buffer (50 mM HEPES, 1 mM EGTA, 1 mM MgCl_2_, 300 mM KCl, 10% glycerol, 0.05% NP-40, 1 mM dithiothreitol, 10 µg/mL leupeptin/pepstatin/chymostatin, pH 7.4). After a final rinse in wash buffer without detergent, bound protein was eluted with 100 mM glycine pH 2.6. Eluted proteins were precipitated by addition of 1/5^th^ volume trichloroacetic acid at 4°C overnight. Precipitated proteins were reduced with TCEP, alkylated with iodoacetamide, and digested with mass-spectrometry grade Lys-C and trypsin (Promega) using S-Trap (Protifi) according to the manufacturer’s instructions. Peptides were separated by liquid chromatography and analyzed on an Orbitrap Elite mass spectrometer (Thermo Fisher). Data were analyzed using Proteome Discoverer Software (Thermo Fisher).

### Western Blotting

Cells expressing individual sgRNAs were induced in 1 μg/mL doxycycline for 3 days before lysis in Laemmli buffer and incubation at 95°C for 5 min. Samples were separated by SDS-PAGE and semi-dry transferred to nitrocellulose. Membranes were blocked for 30 min in blocking buffer (5% BSA in TBS with 0.1% Tween-20) before incubation with anti-phospho-H2A.X (Ser139) antibody (Millipore clone JBW301) at 1:1000 dilution followed by HRP-conjugated secondary antibody (Kindle Biosciences) at 1:5000 dilution. To detect GAPDH as a loading control, HRP-conjugated antibody (ab185059) was applied at 1:20,000 dilution. Membranes were imaged with a KwikQuant Imager (Kindle Biosciences) and quantified using Image Studio software (LI-COR).

### Arrayed imaging experiments with inducible knockout cell lines

Inducible knockout cell lines for immunofluorescence were seeded on poly-L-lysine (Sigma-Aldrich) coated coverslips and fixed in PHEM with 4% formaldehyde for 10 min at 37°C (microtubule staining) or ice cold methanol. Coverslips were washed with PBS, permeabilized with 0.2% Triton X-100 in PBS, and blocked in Abdil buffer (20 mM Tris-HCl, 150 mM NaCl, 0.1% Triton X-100, 3% bovine serum albumin, 0.1% NaN_3_, pH 7.5). Anti-alpha-tubulin (DM1A, Sigma; 1:3000 dilution), anti-CENP-A (Clone 3-19, Invitrogen; 1:1000 dilution) and anti-“Bonsai”/NDC80 (*68*; 1 µg/mL) antibodies in Abdil buffer were used for primary staining. Cy2- and Cy5-conjugated secondary antibodies (Jackson ImmunoResearch Laboratories) were diluted 1:500 with 1 µg/mL Hoechst-33342 (Sigma-Aldrich) in Abdil for subsequent staining. Slides were mounted with ProLong Gold Antifade (Invitrogen) prior to imaging using the microscope configuration described above.

For quantifications of Ndc80 and CENP-A kinetochore stain intensity, sections of cells were maximum intensity projected and cropped in Fiji (*69*). Integrated fluorescence intensity of mitotic kinetochores was measured with a custom pipeline in CellProfiler (*70*). The median intensity of a 5-pixel wide region surrounding each kinetochore was used to background subtract each measurement.

For time lapse analysis of individual knockout cell lines, cells were induced in 12-well polymer-bottomed plates (Cellvis) with 1 μg/mL doxycycline for 3 days, refreshing doxycycline media each day. On day 3 or 4 post-Cas9 induction, cells were moved to CO_2_-independent media (Gibco) supplemented with 10% FBS, 100 U/mL penicillin and streptomycin, and 2 mM L-glutamine before imaging using the microscope configuration described above.

### RNA-sequencing of inducible knockout cell lines

Inducible knockout cells were seeded in 1 μg/mL doxycycline, and doxycycline media was refreshed each day for 3 days before harvest of a mitotically-enriched cell population by shake-off. Cells were washed in PBS before snap-freezing pellets of 100,000 cells in liquid nitrogen. Cells were lysed with 1X TCL buffer (Qiagen) supplemented with 1% beta-mercaptoethanol. Smart-seq2 was performed as described previously (*71*) to prepare libraries for sequencing on an Illumina NovaSeq or MiniSeq. Transcript counts were quantified using kallisto (*72*) with default parameters and Gencode release 21. Differential expression analysis was performed using edgeR (*73*) with default settings; comparisons to control samples were completed using the exactTest function. The Benjamini-Hochberg procedure was used to estimate FDR to identify differentially expressed genes (FDR < 0.05, effect size > 2-fold change).

**Fig. S1.**
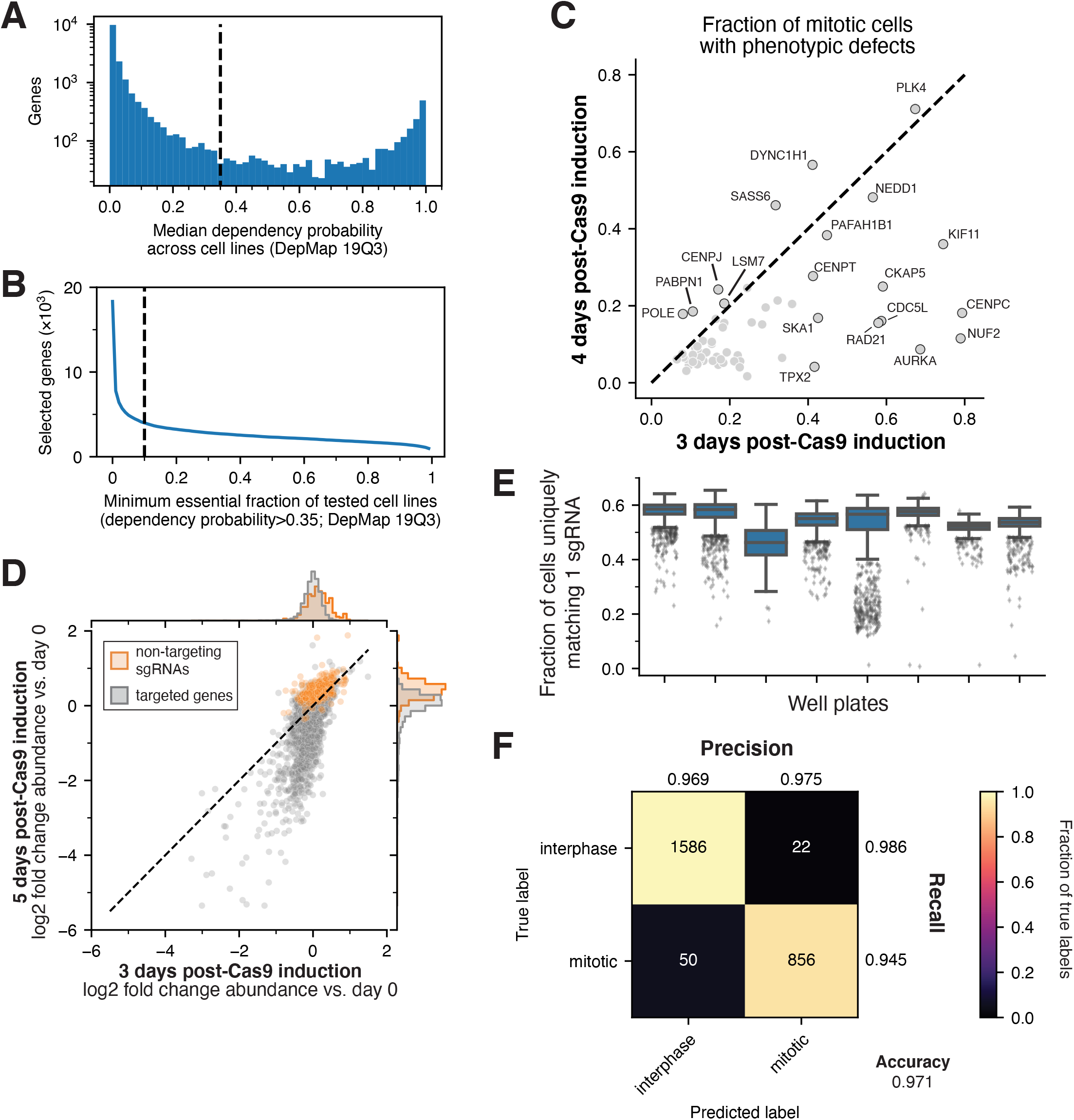
Optimization of image-based pooled screening for the definition of essential gene function. (**A**) Histogram of median dependency probability across cell lines in the DepMap dataset, indicating the threshold chosen (0.35) for defining individual essential cell lines for each gene in (B). (**B**) Number of genes identified as essential for at least the specified fraction of tested cell lines in DepMap. Genes that were essential in at least 10% of tested cell lines were selected for the screen, in addition to those selected from additional sources (see Methods). (**C**) Scatter plot showing the results from small-scale screens of 400 gene targets. This compares the fraction of mitotic cells with phenotypic defects for established cell division factors at 3 and 4 days post-Cas9 induction. Overall, mitotic phenotypes were more commonly observed at the earlier time point. (**D**) Scatter plot showing mean change in abundance within the 20,445 sgRNA primary screen library at 3 and 5 days post-Cas9 induction, each time point relative to pre-induction (day 0). N=2 screen replicates were performed, averaged across sgRNAs targeting the same gene. Orange indicates non-targeting control sgRNAs. Many gene targets begin to drop out of the population at day 5 due to fitness defects. Based on this data and from (C), 78 hours post-Cas9 induction was chosen as the fixation time point for our image-based screen to maximize observable phenotypes. (**E**) Boxplot demonstrating *in situ* sequencing quality in our fixed-cell image-based pooled screen. Sequencing quality was consistent across the eight imaging plates, with the majority of imaging tiles exceeding 50% of cells with sequencing reads that uniquely match a single sgRNA sequence from the library. N = 1,665 or 1,998 imaging tiles in each plate column. (**F**) Confusion matrix demonstrating performance of the support vector classifier in distinguishing interphase and mitotic cells, 5-fold cross-validation with N=2,514 manually annotated cell images.

**Fig. S2.**
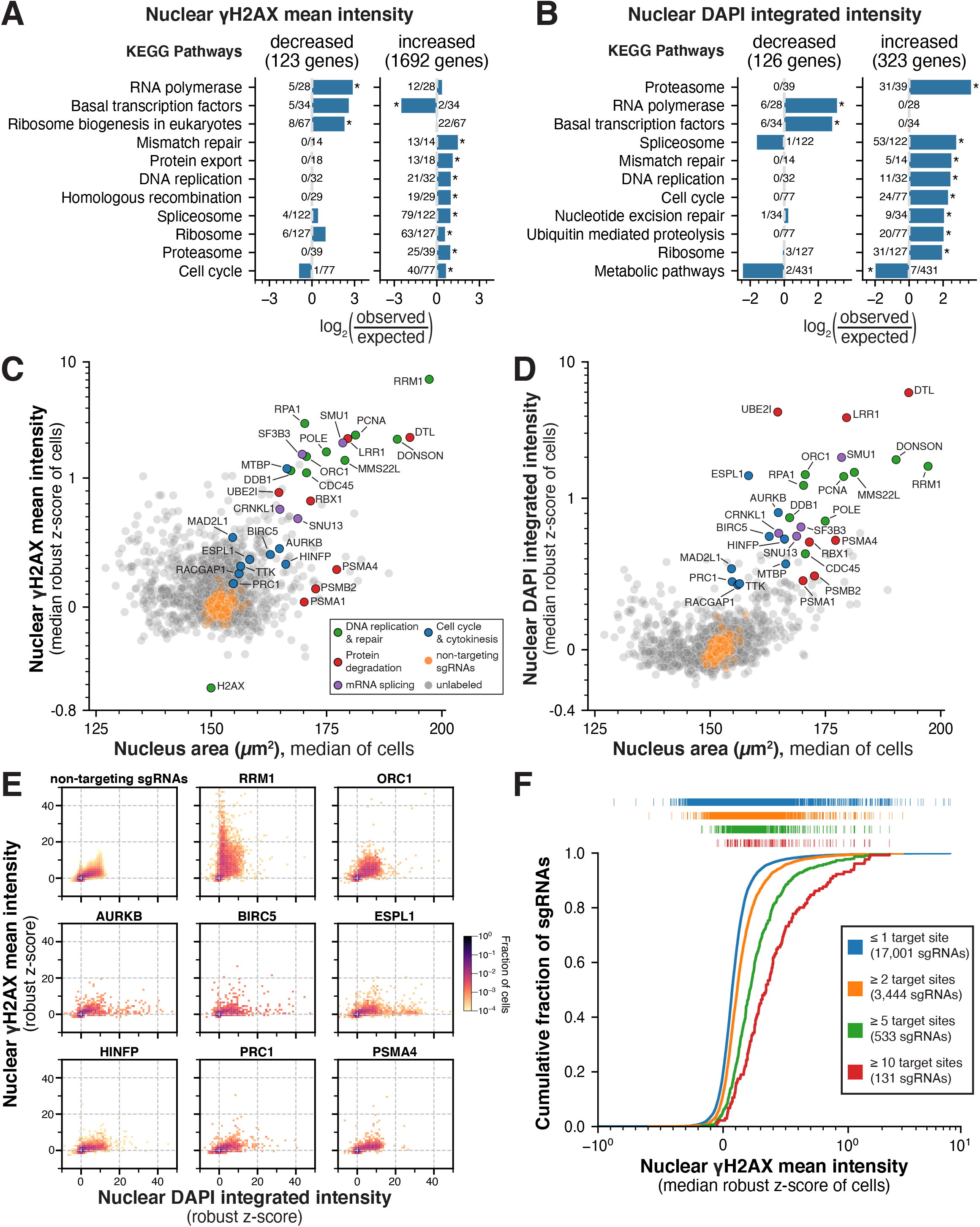
Analysis of interphase nuclear phenotypes. (**A**) Bar graph indicating over-representation of KEGG pathways among gene targets exhibiting decreased or increased nuclear γH2AX mean intensity. *FDR<0.05 (**B**) Bar graph of over-representation analysis results as in (A) among gene targets with decreased or increased nuclear DNA (DAPI) integrated intensity. *FDR<0.05 (**C**) Scatter plot showing summary gene scores (see Methods) for mean nuclear γH2AX intensity compared to nuclear area, showing a subset of gene knockouts with increases in both γH2AX and nuclear area. Summary γH2AX scores are plotted on a symmetric log scale (linear between -1 and 1) and labeled genes are colored by functional category. (**D**) Scatter plot showing summary gene scores for integrated nuclear DNA (DAPI) intensity compared to nuclear area as in (C). DNA content is relatively constant across gene targets exhibiting a range of cell areas, although a subset demonstrates increased nuclear area and DNA. Summary DAPI scores are plotted on a symmetric log scale (linear between -1 and 1) and labeled genes are colored by functional category. (**E**) Bivariate histograms of integrated nuclear DNA intensity and mean nuclear γH2AX intensity, displaying single-cell distributions for all cells expressing non-targeting sgRNAs (top left) and selected gene targets. Knockouts of genes that regulate chromosome segregation or cytokinesis result in more cells with increased DNA content, but only modest increases in γH2AX intensity. Histogram bins containing a fraction of total cells for a given gene target that is less than 1 in 104 cells are not displayed. (**F**) Cumulative distribution plots of mean nuclear γH2AX intensity (DNA damage phenotype) for subsets of sgRNAs sequences with increasing numbers of genomic target sites. Non-targeting control sgRNAs and sgRNAs targeting a single genomic locus (blue) include the vast majority of sgRNAs in the library and displayed minimal DNA damage on average in the screen. In contrast, sgRNAs with increasing numbers of target sites (orange, green, and red) displayed distribution shifts toward increasing DNA damage. Tick marks above the plot indicate individual sgRNA phenotype scores, which are the median robust z-scores measured across all cells with a given sgRNA. Genomic target sites are the number of cutting frequency determination (CFD) bin 1 matches (see *19*).

**Fig. S3.**
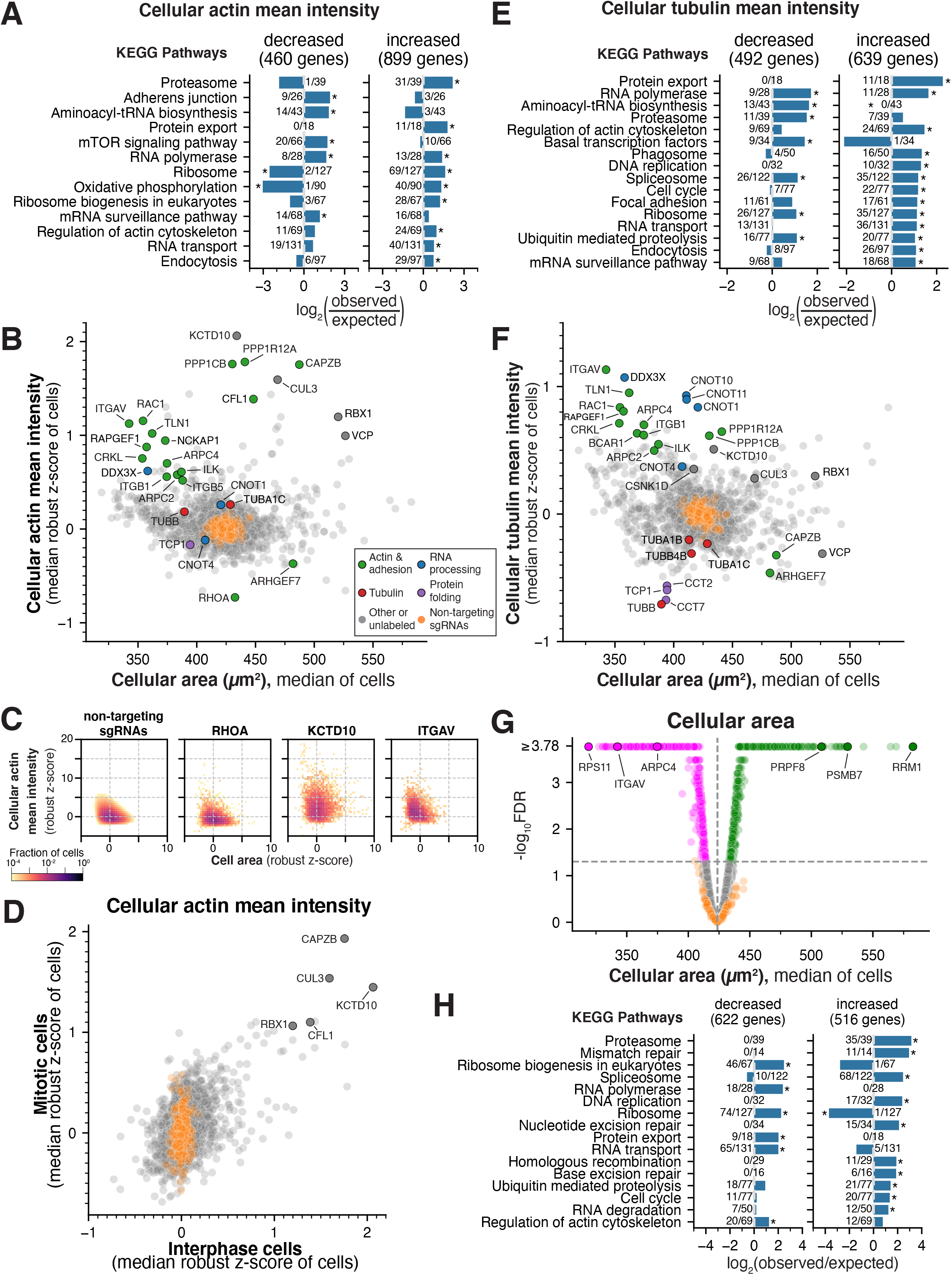
Analysis of interphase cytoskeletal phenotypes. (**A**) Bar graph indicating over-representation of KEGG pathways among gene targets with decreased or increased mean cellular actin (phalloidin) intensity in interphase cells. *FDR<0.05 (**B**) Scatter plot indicating summary gene scores (see Methods) for mean cellular actin intensity compared to cell area. A subset of gene knockouts display increased actin staining together with decreased cell area due to disrupted cellular adhesion. Labeled genes are colored by functional category. (**C**) Bivariate histograms of mean cellular actin intensity and cellular area, displaying single-cell distributions for all cells expressing non-targeting sgRNAs (left) and selected gene targets. Knockouts of genes that regulate cellular adhesion (e.g., ITGAV) show a distribution of cells shifted toward lower cellular area and correspondingly increased mean actin intensity. Histogram bins containing a fraction of total cells for a given gene target that is less than 1 in 104 cells are not displayed. (**D**) Scatter plot comparing mean cellular actin intensity summary scores between interphase and mitotic cell populations, indicating factors that robustly affect actin structures throughout the cell cycle. (**E**) Bar graph of over-representation analysis results as in (A) for gene targets demonstrating decreased or increased mean cellular tubulin intensity. *FDR<0.05 (**F**) Scatter plot showing summary gene scores for mean cellular tubulin intensity compared to cell area as in (B). Similar to actin, a subset of gene knockouts display increased tubulin staining in combination with decreased cell area. (**G**) Volcano plot for interphase cell area across gene targets in the screen, showing a wide range of decreased (magenta) and increased (green) cell areas (FDR<0.05). Raw P-values were computed by comparing gene targets to a bootstrapped null distribution of cells expressing non-targeting sgRNAs (see Methods), with false discovery rate (FDR) estimated using the Benjamini-Hochberg procedure. (**H**) Bar graph of over-representation analysis results as in (A) for gene targets that result in decreased or increased cell area.

**Fig. S4.**
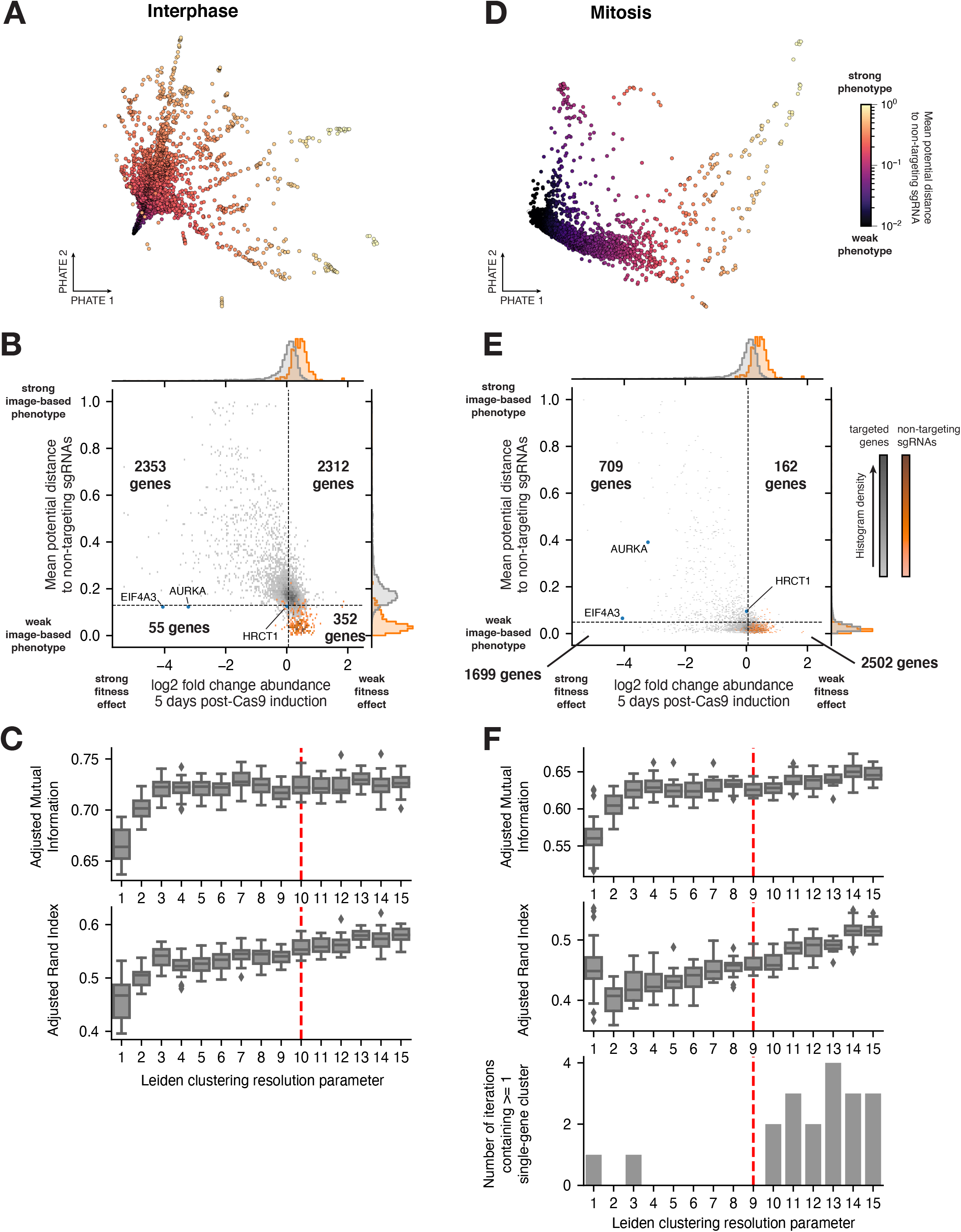
PHATE analysis of multi-dimensional phenotypes. (**A**) Two-dimensional representation of the interphase phenotype landscape of gene targets in the primary screen using PHATE (35; see Methods). Data points are colored by the mean potential distance to non-targeting control sgRNAs computed by PHATE for the interphase phenotype profiles, normalized between 0 and 1. Gene targets with increasing phenotype strength intuitively radiate outward from a dense region containing non-targeting sgRNAs. (**B**) Bivariate histogram showing the joint distribution of image-based interphase phenotype strength from (A) together with the strength of knockout fitness effect in the screening cell line (mean change in sgRNA abundance within the library after 5 days of Cas9 induction, n=2 screen replicates averaged across sgRNAs targeting the same gene). >90% of gene targets exhibit a measurable interphase phenotype in the image-based screen (potential distance greater than the 95th percentile of non-targeting sgRNAs). Of the remaining 407 genes, only 55 demonstrate a meaningful fitness effect in the tested cell line (log2 fold change abundance less than the 5th percentile of non-targeting sgRNAs). Labeled genes are those that display a fitness effect and no interphase phenotype, but do show a measurable mitotic phenotype in (E). (**C**) Boxplot illustrating selection of the Leiden clustering (64) resolution parameter for interphase phenotypes. For 20 repetitions at each resolution, 90% of gene targets were sampled without replacement and clustered using PHATE and Leiden algorithms in series. Each sampled cluster solution was then compared to the full dataset clusters using adjusted mutual information (top) and adjusted Rand index (bottom). The dotted line indicates the chosen resolution, selected based on the plateau in robustness of clustering solutions. (**D**) Two dimensional representation of the mitotic phenotype landscape of gene targets as in (A), colored by the mean potential distance to non-targeting control sgRNAs computed by PHATE for the mitotic phenotype profiles. (**E**) Bivariate histogram showing the joint distribution of image-based mitotic phenotype strength from (D) together with the strength of knockout fitness effect in the screening cell line as in (B). The threshold for measurable mitotic phenotypes is the 95th percentile of potential distance among non-targeting control sgRNAs. Labeled genes indicate those that display a fitness effect and mitotic phenotype, but do not exhibit an interphase phenotype in (B). (**F**) Boxplot illustrating selection of the Leiden clustering resolution parameter for mitotic phenotypes using the same procedure as (C). A resolution parameter of 9 was chosen in part due to the increased presence of single-gene clusters with resolution ≥10 (bottom).

**Fig. S5.**
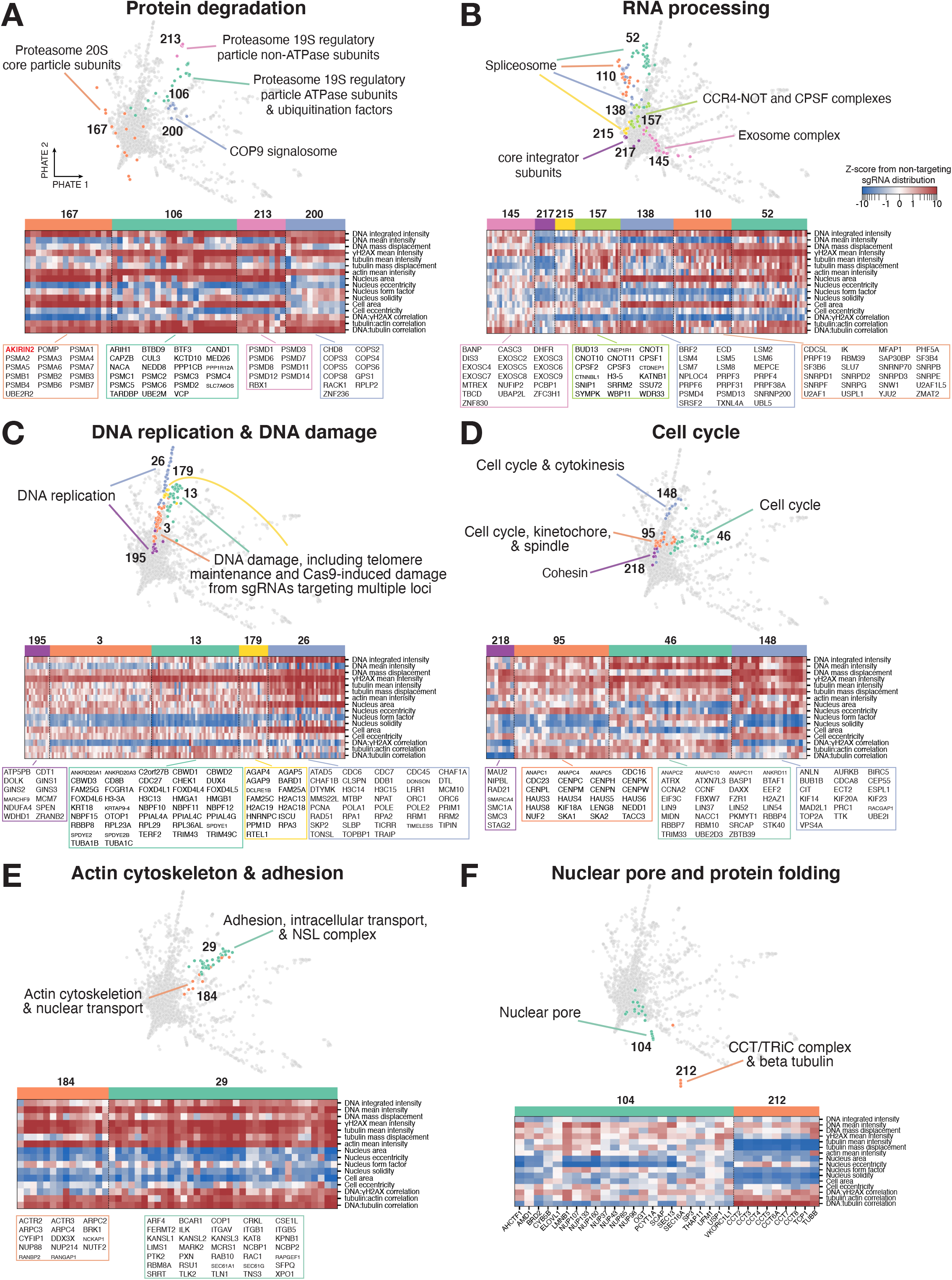
Interphase phenotypes enable detailed clustering of specific functional categories. Two-dimensional PHATE representations of interphase phenotype clusters and corresponding heat maps of a manually-selected subset of specific parameters for a broad range of functional categories, as in Fig. 3. Distinct and coherent phenotypes are observable for genes involved in processes such as (**A**) protein degradation (including the recently-characterized gene AKIRIN2; (**B**) RNA processing; (**C**) DNA replication and DNA damage; (**D**) cell cycle function; (**E**) actin cytoskeleton and cellular adhesion; and (**F**) nuclear pore function and protein folding. Numbers indicate individual interphase cluster identities. All genes from selected clusters are listed below each heatmap. Parameters are presented as z-scores from the distribution of non-targeting sgRNAs, visualized on a symmetric log scale (linear between -1 and 1).

**Fig. S6.**
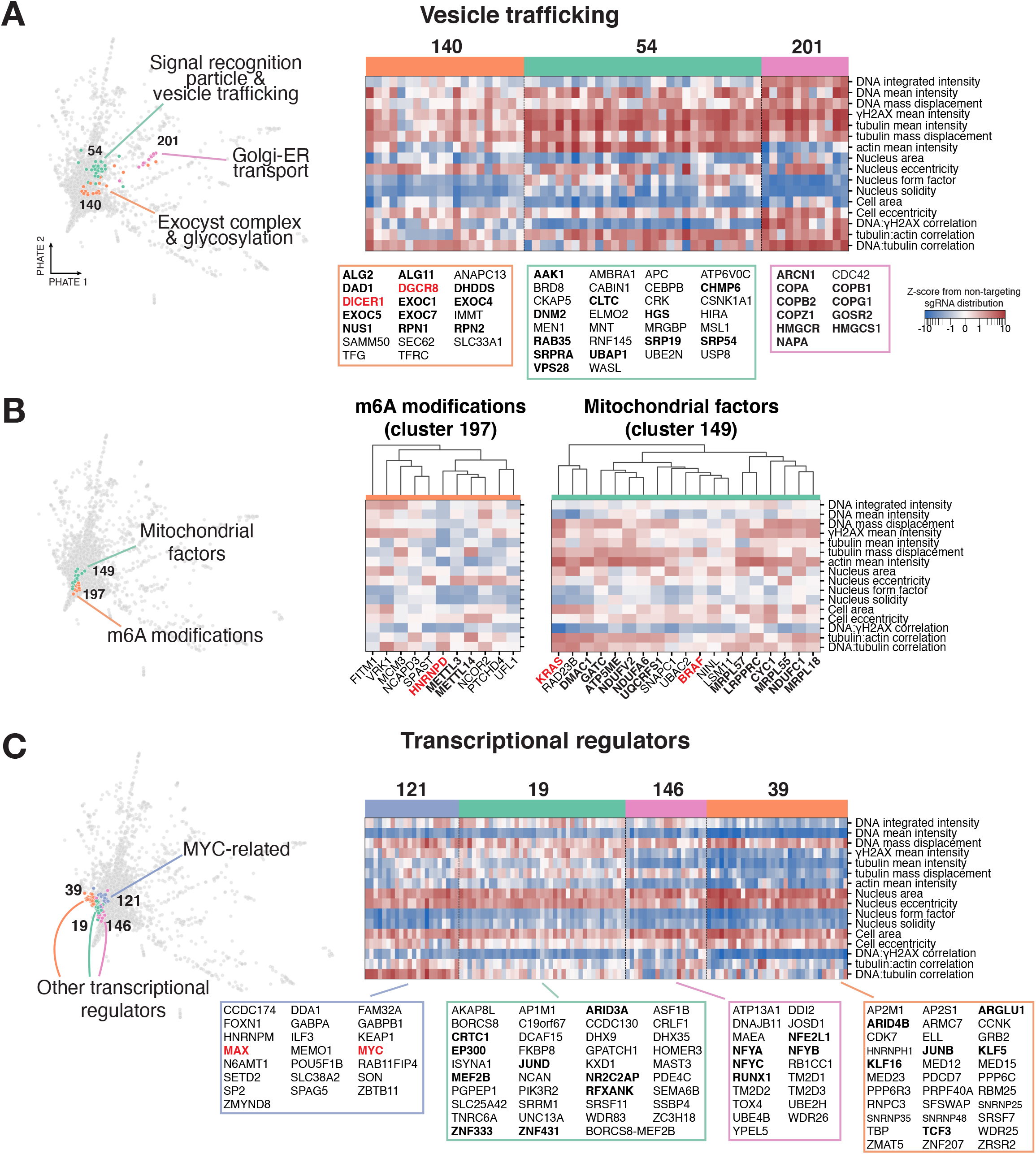
Interphase cluster analysis reveals novel functional associations for established factors. Two-dimensional PHATE representations of interphase phenotype clusters and corresponding heat maps of a manually-selected subset of specific parameters, as in Fig. 3. (**A**) Gene targets involved in vesicle trafficking and related processes exhibit distinct image-based phenotypes, despite the absence of membrane-targeted phenotype stains in the screen. Targeting DICER1 and the associated factor DGCR8 unexpectedly results in a phenotype similar to exocyst complex knockouts. (**B**) Additional clusters identify a co-functional role of HNRNPD and m6A modifications, as well as a relationship between mitochondrial function and KRAS/BRAF signaling. (**C**) Transcriptional regulators show interrelated phenotypes, with an apparent unique phenotype for cluster 121 containing MYC and MAX, suggesting the presence of additional MYC-associated factors.

**Fig. S7.**
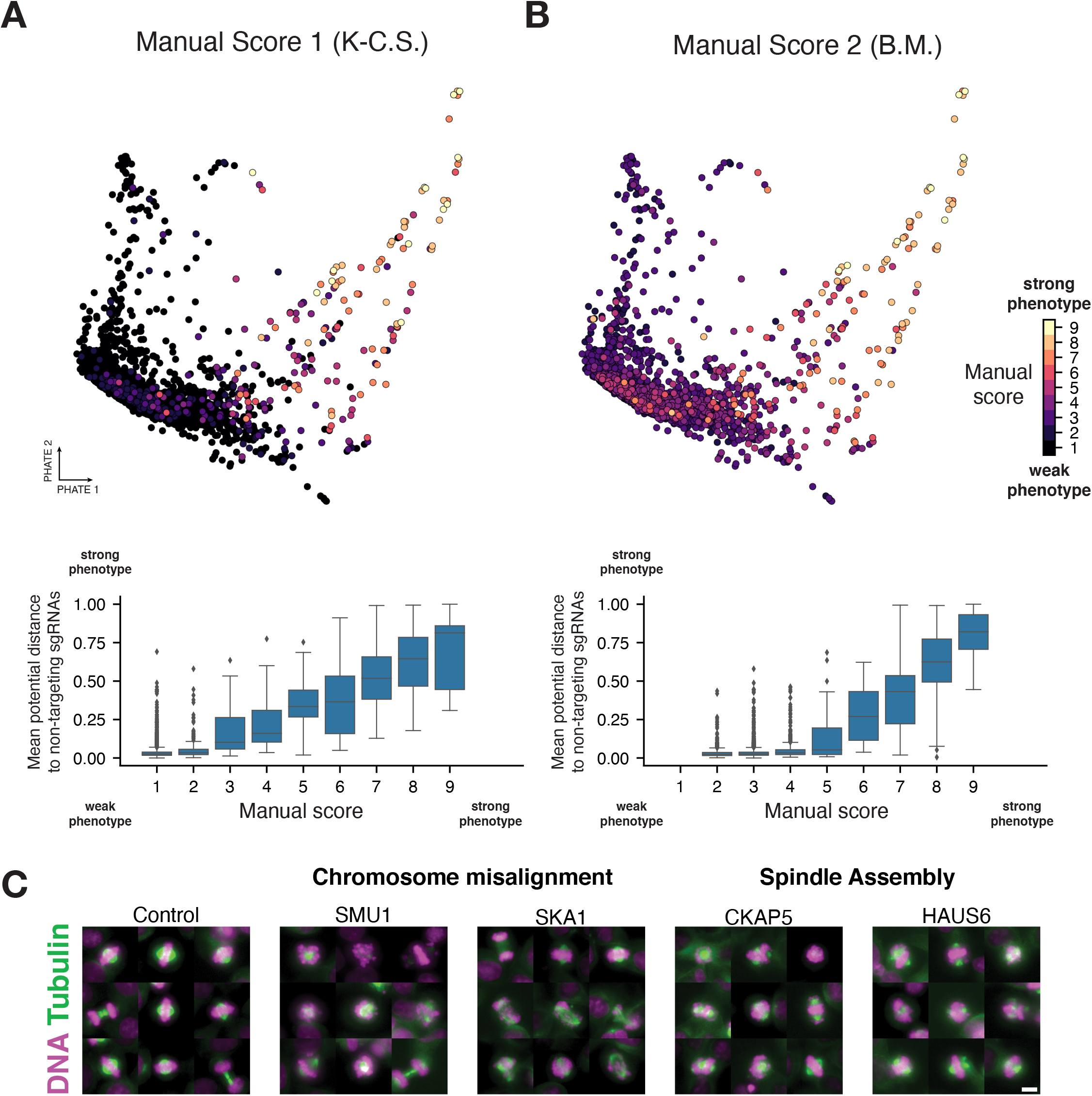
Computational and manual scoring show strong agreement for mitotic phenotypes. (**A** and **B**, top) Two-dimensional PHATE representations of mitotic phenotypes, colored by manual phenotype severity scores independently assigned by two individuals using anonymized gene names. (A and B, bottom) Box plots demonstrating strong agreement between manual phenotype scores and computational phenotype strength (mean potential distance to non-targeting sgRNAs from mitotic PHATE analysis, normalized between 0 and 1. (**C**) Example images of mitotic cells from the primary screen visualizing DNA (magenta) and microtubules (green) from gene targets selected in the computational and/or visual analysis highlight the diversity of identified mitotic phenotypes (see also Fig. 4D). Control images are reproduced from Fig. 4D. Scale bar, 10 µm.

**Fig. S8.**
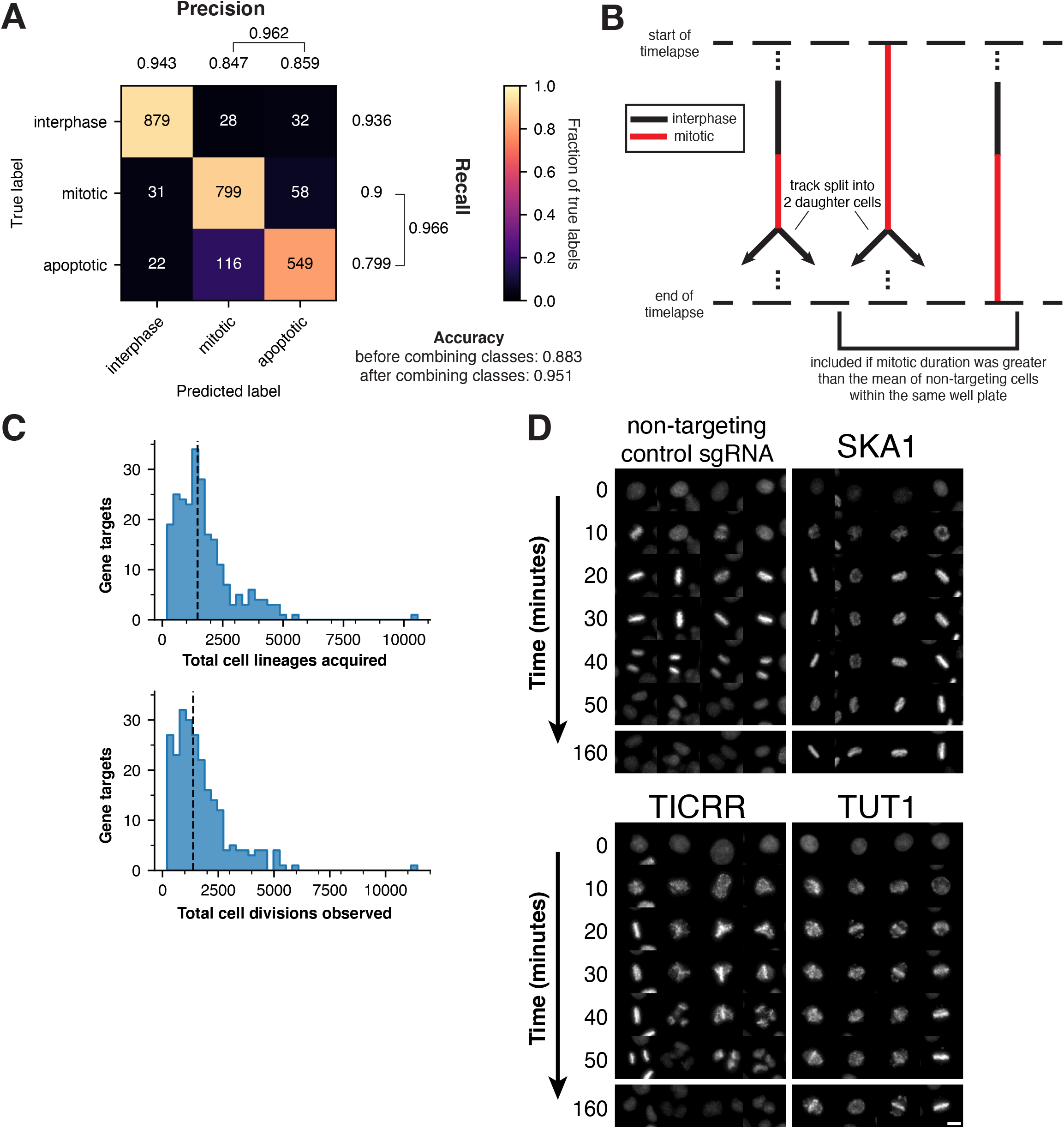
Optical pooled screening for live-cell mitotic phenotypes. (**A**) Confusion matrix demonstrating performance of the support vector classifier in distinguishing interphase, mitotic, and apoptotic cells from the live-cell screen. 5-fold cross-validation with n=2,514 manually annotated cell images. Due to the relative difficulty of differentiating mitotic and apoptotic cells from H2B-mCherry fluorescence alone, the mitotic and apoptotic classes were combined after inference (precision and recall after combining classes indicated by brackets). (**B**) Criteria for identifying a cell division event in the live-cell screen analysis. Cell lineages that were not tracked across the entire time course were excluded. (**C**) Histograms of the total cell lineages acquired (top) and cell divisions observed (bottom) across both day 3 and day 4 time courses for each gene target. (**D**) Example images of H2B-mCherry fluorescence from the live-cell screen at the indicated time points after mitotic entry for selected gene targets. Each displayed knockout demonstrates increased mitotic duration and chromosome alignment defects relative to the non-targeting control sgRNA. Control images are reproduced from Fig. 5B. Scale bar, 10 µm.

**Fig. S9.**
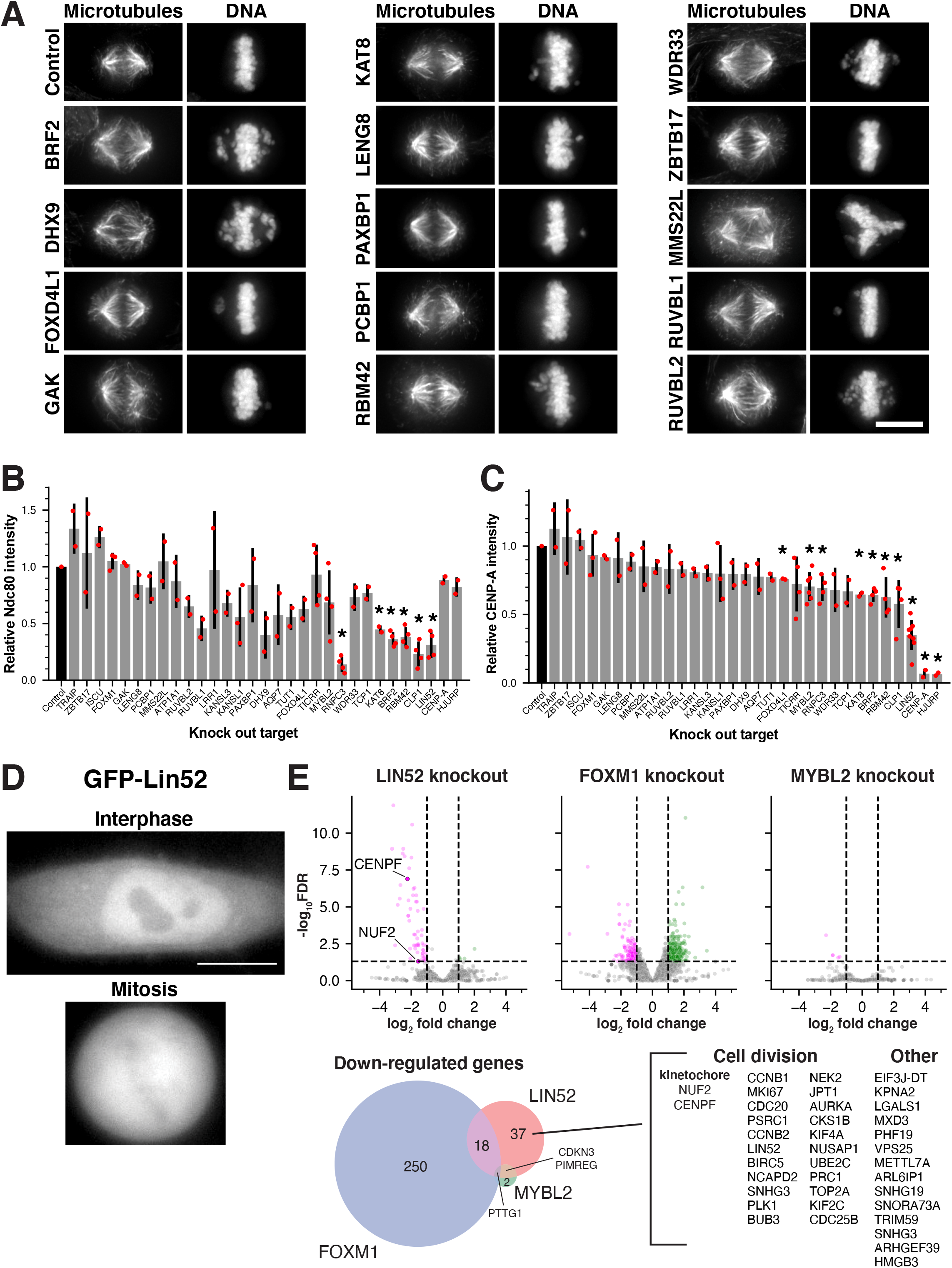
Targeted analysis of mitotic phenotypes. (**A**) Immunofluorescence images of individual cell lines stably expressing a single sgRNA targeting each gene of interest to confirm live-cell pooled screen phenotypes and enable visualization at higher resolution across a single population. Images are deconvolved maximum intensity projections of fixed cells stained for microtubules (anti-alpha-tubulin) and DNA (Hoechst). Scale bar, 10 µm. (**B**) Bar plot showing kinetochore-localized intensity of the outer kinetochore microtubule-binding protein Ndc80 in inducible knockout cell lines for all 29 genes pursued from the live-cell screen, along with CENP-A and HJURP controls. Each data point represents the median kinetochore signal of one experiment for >10 cells per gene target. Values are normalized relative to negative control cells from the same experiment. *P<0.05 by two-sided independent T-test relative to negative control cells, Bonferroni-adjusted. (**C**) Bar plot of kinetochore-localized intensity for the inner kinetochore centromere-specific histone CENP-A in inducible knockout cell lines; experiment design as in (B). *P<0.01 by two-tailed independent T-test relative to control cells. (**D**) Fluorescence images of human cells expressing GFP-LIN52, indicating LIN52 nuclear localization in interphase cells and non-specific localization in mitotic cells. Scale bar, 10 µm. (**E**) RNA-seq analysis of DREAM complex-related inducible knockout cell lines in a mitotically-enriched population (see Methods). Targeting LIN52 and FOXM1 resulted in many differentially expressed genes compared to cells expressing the HS1 control sgRNA, while MYBL2 knockouts displayed minimal transcriptional effects in the tested cell population (top). Disruption of LIN52 resulted in 37 unique down-regulated genes relative to FOXM1 and MYBL2, including several genes involved in cell division and the kinetochore components NUF2 and CENPF. However, the down-regulation of these genes alone is unlikely to explain the strong effect of LIN52 knockout on both CENP-A and Ndc80 localization.

## OTHER SUPPLEMENTAL MATERIALS

**Movie S1**. Example time lapse montages from the secondary live cell screen.

**Movie S2**. Example time lapse images of individual knockout cell lines.

**Movie S3**. Time lapse movies showing Lin52 complex knockout phenotypes.

**Data S1**. CRISPR sgRNA sequences used in this study.

**Data S2**. Primary screen results and extracted image features.

**Data S3**. Secondary screen results and extracted image features.

**Data S4**. Cell lines used in this study.

## REFERENCES

1. T. Wang, J. J. Wei, D. M. Sabatini, E. S. Lander, Genetic Screens in Human Cells Using the CRISPR-Cas9 System. Science. 343, 80–84 (2014).

2. K. J. Condon, J. M. Orozco, C. H. Adelmann, J. B. Spinelli, P. W. van der Helm, J. M. Roberts, T. Kunchok, D. M. Sabatini, Genome-wide CRISPR screens reveal multitiered mechanisms through which mTORC1 senses mitochondrial dysfunction. Proc. Natl. Acad. Sci. U. S. A. 118, e2022120118 (2021).

3. J. Nieuwenhuis, A. Adamopoulos, O. B. Bleijerveld, A. Mazouzi, E. Stickel, P. Celie, M. Altelaar, P. Knipscheer, A. Perrakis, V. A. Blomen, T. R. Brummelkamp, Vasohibins encode tubulin detyrosinating activity. Science. 358, 1453– 1456 (2017).

4. A. Dixit, O. Parnas, B. Li, J. Chen, C. P. Fulco, L. Jerby-Arnon, N. D. Marjanovic, D. Dionne, T. Burks, R. Raychowdhury, B. Adamson, T. M. Norman, E. S. Lander, J. S. Weissman, N. Friedman, A. Regev, Perturb-Seq: Dissecting Molecular Circuits with Scalable Single-Cell RNA Profiling of Pooled Genetic Screens. Cell. 167, 1853-1866. e17 (2016).

5. G. Kanfer, S. A. Sarraf, Y. Maman, H. Baldwin, E. Dominguez-Martin, K. R. Johnson, M. E. Ward, M. Kampmann, J. Lippincott-Schwartz, R. J. Youle, Image-based pooled whole-genome CRISPRi screening for subcellular phenotypes. J. Cell Biol. 220, e202006180 (2021).

6. X. Yan, N. Stuurman, S. A. Ribeiro, M. E. Tanenbaum, M. A. Horlbeck, C. R. Liem, M. Jost, J. S. Weissman, R. D. Vale, High-content imaging-based pooled CRISPR screens in mammalian cells. J. Cell Biol. 220, e202008158 (2021).

7. N. Hasle, A. Cooke, S. Srivatsan, H. Huang, J. J. Stephany, Z. Krieger, D. Jackson, W. Tang, S. Pendyala, R. J. Monnat Jr., C. Trapnell, E. M. Hatch, D. M. Fowler, High-throughput, microscope-based sorting to dissect cellular heterogeneity. Mol. Syst. Biol. 16, e9442 (2020).

8. D. Feldman, A. Singh, J. L. Schmid-Burgk, R. J. Carlson, A. Mezger, A. J. Garrity, F. Zhang, P. C. Blainey, Optical Pooled Screens in Human Cells. Cell. 179, 787-799. e17 (2019).

9. D. Feldman, L. Funk, A. Le, R. J. Carlson, M. Leiken, F. Tsai, B. Soong, A. Singh, P. C. Blainey, Pooled genetic perturbation screens with image-based phenotypes. Nat. Protoc. (In press).

10. V. A. Blomen, P. Májek, L. T. Jae, J. W. Bigenzahn, J. Nieuwenhuis, J. Staring, R. Sacco, F. R. van Diemen, N. Olk, A. Stukalov, C. Marceau, H. Janssen, J. E. Carette, K. L. Bennett, J. Colinge, G. Superti-Furga, T. R. Brummelkamp, Gene essentiality and synthetic lethality in haploid human cells. Science. 350, 1092–1096 (2015).

11. J. M. Dempster, J. Rossen, M. Kazachkova, J. Pan, G. Kugener, D. E. Root, A. Tsherniak, Extracting Biological Insights from the Project Achilles Genome-Scale CRISPR Screens in Cancer Cell Lines. bioRxiv, 720243 (2019).

12. DepMap, Broad., DepMap 19Q3 Public. figshare. Dataset doi:10.6084/m9.figshare.9201770.v2.

13. T. Hart, M. Chandrashekhar, M. Aregger, Z. Steinhart, K. R. Brown, G. MacLeod, M. Mis, M. Zimmermann, A. Fradet-Turcotte, S. Sun, P. Mero, P. Dirks, S. Sidhu, F. P. Roth, O. S. Rissland, D. Durocher, S. Angers, J. Moffat, High-Resolution CRISPR Screens Reveal Fitness Genes and Genotype-Specific Cancer Liabilities. Cell. 163, 1515–1526 (2015).

14. M. A. Horlbeck, L. A. Gilbert, J. E. Villalta, B. Adamson, R. A. Pak, Y. Chen, A. P. Fields, C. Y. Park, J. E. Corn, M. Kampmann, J. S. Weissman, Compact and highly active next-generation libraries for CRISPR-mediated gene repression and activation. eLife. 5, e19760 (2016).

15. R. M. Meyers, J. G. Bryan, J. M. McFarland, B. A. Weir, A. E. Sizemore, H. Xu, N. V. Dharia, P. G. Montgomery, G. S. Cowley, S. Pantel, A. Goodale, Y. Lee, L. D. Ali, G. Jiang, R. Lubonja, W. F. Harrington, M. Strickland, T. Wu, D. C. Hawes, V. A. Zhivich, M. R. Wyatt, Z. Kalani, J. J. Chang, M. Okamoto, K. Stegmaier, T. R. Golub, J. S. Boehm, F. Vazquez, D. E. Root, W. C. Hahn, A. Tsherniak, Computational correction of copy number effect improves specificity of CRISPR–Cas9 essentiality screens in cancer cells. Nat. Genet. 49, 1779– 1784 (2017).

16. K. Tzelepis, H. Koike-Yusa, E. De Braekeleer, Y. Li, E. Metzakopian, O. M. Dovey, A. Mupo, V. Grinkevich, M. Li, M. Mazan, M. Gozdecka, S. Ohnishi, J. Cooper, M. Patel, T. McKerrell, B. Chen, A. F. Domingues, P. Gallipoli, S. Teichmann, H. Ponstingl, U. McDermott, J. Saez-Rodriguez, B. J. P. Huntly, F. Iorio, C. Pina, G. S. Vassiliou, K. Yusa, A CRISPR Dropout Screen Identifies Genetic Vulnerabilities and Therapeutic Targets in Acute Myeloid Leukemia. Cell Rep. 17, 1193–1205 (2016).

17. T. Wang, K. Birsoy, N. W. Hughes, K. M. Krupczak, Y. Post, J. J. Wei, E. S. Lander, D. M. Sabatini, Identification and characterization of essential genes in the human genome. Science. 350, 1096–1101 (2015).

18. T. Wang, H. Yu, N. W. Hughes, B. Liu, A. Kendirli, K. Klein, W. W. Chen, E. S. Lander, D. M. Sabatini, Gene Essentiality Profiling Reveals Gene Networks and Synthetic Lethal Interactions with Oncogenic Ras. Cell. 168, 890-903. e15 (2017).

19. J. G. Doench, N. Fusi, M. Sullender, M. Hegde, E. W. Vaimberg, K. F. Donovan, I. Smith, Z. Tothova, C. Wilen, R. Orchard, H. W. Virgin, J. Listgarten, D. E. Root, Optimized sgRNA design to maximize activity and minimize off-target effects of CRISPR-Cas9. Nat. Biotechnol. 34, 184–191 (2016).

20. T. Hart, A. H. Y. Tong, K. Chan, J. Van Leeuwen, A. Seetharaman, M. Aregger, M. Chandrashekhar, N. Hustedt, S. Seth, A. Noonan, A. Habsid, O. Sizova, L. Nedyalkova, R. Climie, L. Tworzyanski, K. Lawson, M. A. Sartori, S. Alibeh, D. Tieu, S. Masud, P. Mero, A. Weiss, K. R. Brown, M. Usaj, M. Billmann, M. Rahman, M. Constanzo, C. L. Myers, B. J. Andrews, C. Boone, D. Durocher, J. Moffat, Evaluation and Design of Genome-Wide CRISPR/SpCas9 Knockout Screens. G3 Bethesda Md. 7, 2719–2727 (2017).

21. K. L. McKinley, I. M. Cheeseman, Large-Scale Analysis of CRISPR/Cas9 Cell-Cycle Knockouts Reveals the Diversity of p53-Dependent Responses to Cell-Cycle Defects. Dev. Cell. 40, 405-420.e2 (2017).

22. Y. Dang, G. Jia, J. Choi, H. Ma, E. Anaya, C. Ye, P. Shankar, H. Wu, Optimizing sgRNA structure to improve CRISPR-Cas9 knockout efficiency. Genome Biol. 16, 280 (2015).

23. P. Datlinger, A. F. Rendeiro, C. Schmidl, T. Krausgruber, P. Traxler, J. Klughammer, L. C. Schuster, A. Kuchler, D. Alpar, C. Bock, Pooled CRISPR screening with single-cell transcriptome readout. Nat. Methods. 14, 297–301 (2017).

24. A. Sancar, L. A. Lindsey-Boltz, K. Unsal-Kaçmaz, S. Linn, Molecular mechanisms of mammalian DNA repair and the DNA damage checkpoints. Annu. Rev. Biochem. 73, 39– 85 (2004).

25. C. Pederiva, S. Böhm, A. Julner, M. Farnebo, Splicing controls the ubiquitin response during DNA double-strand break repair. Cell Death Differ. 23, 1648–1657 (2016).

26. R. Shi, W. Hou, Z.-Q. Wang, X. Xu, Biogenesis of Iron– Sulfur Clusters and Their Role in DNA Metabolism. Front. Cell Dev. Biol. 9, 2676 (2021).

27. F. Villa, R. Fujisawa, J. Ainsworth, K. Nishimura, M. Lie-A-Ling, G. Lacaud, K.P. Labib, CUL2LRR1, TRAIP and p97 control CMG helicase disassembly in the mammalian cell cycle. EMBO Rep. 22, e52164 (2021).

28. T. D. Pollard, Actin and Actin-Binding Proteins. Cold Spring Harb. Perspect. Biol. 8, a018226 (2016).

29. H. V. Goodson, E. M. Jonasson, Microtubules and Microtubule-Associated Proteins. Cold Spring Harb. Perspect. Biol. 10, a022608 (2018).

30. L. Li, W. Zhang, Y. Liu, X. Liu, L. Cai, J. Kang, Y. Zhang, W. Chen, C. Dong, Y. Zhang, M. Wang, W. Wei, L. Jia, The CRL3BTBD9 E3 ubiquitin ligase complex targets TNFAIP1 for degradation to suppress cancer cell migration. Signal Transduct. Target. Ther. 5, 1–9 (2020).

31. F. Rodríguez-Pérez, A. G. Manford, A. Pogson, A. J. Ingersoll, B. Martínez-González, M. Rape, Ubiquitin-dependent remodeling of the actin cytoskeleton drives cell fusion. Dev. Cell. 56, 588-601.e9 (2021).

32. D. Zyss, H. Ebrahimi, F. Gergely, Casein kinase I delta controls centrosome positioning during T cell activation. J. Cell Biol. 195, 781–797 (2011).

33. M. A. Collart, The Ccr4-Not complex is a key regulator of eukaryotic gene expression. WIREs RNA. 7, 438–454 (2016).

34. H. Cantwell, P. Nurse, Unravelling nuclear size control. Curr. Genet. 65, 1281–1285 (2019).

35. K. R. Moon, D. van Dijk, Z. Wang, S. Gigante, D. B. Burkhardt, W. S. Chen, K. Yim, A. van den Elzen, M. J. Hirn, R. R. Coifman, N. B. Ivanova, G. Wolf, S. Krishnaswamy, Visualizing structure and transitions in high-dimensional biological data. Nat. Biotechnol. 37, 1482–1492 (2019).

36. D. C. I. Goberdhan, C. Wilson, A. L. Harris, Amino Acid Sensing by mTORC1: Intracellular Transporters Mark the Spot. Cell Metab. 23, 580–589 (2016).

37. M.-S. Song, J. J. Rossi, Molecular mechanisms of Dicer: endonuclease and enzymatic activity. Biochem. J. 474, 1603–1618 (2017).

38. Z. Cao, K. A. Budinich, H. Huang, D. Ren, B. Lu, Z. Zhang, Q. Chen, Y. Zhou, Y.-H. Huang, F. Alikarami?, M. C. Kingsley, A. K. Lenard, A. Wakabayashi, E. Khandros, W. Bailis, J. Qi, M. P. Carroll, G. A. Blobel, R. B. Faryabi, K. M. Bernt, S. L. Berger, J. Shi, ZMYND8-regulated IRF8 transcription axis is an acute myeloid leukemia dependency. Mol. Cell. 81, 3604-3622.e10 (2021).

39. S. Singh, A. Vanden Broeck, L. Miller, M. Chaker-Margot, S. Klinge, Nucleolar maturation of the human small subunit processome. Science. 373, eabj5338 (2021).

40. M. de Almeida, M. Hinterndorfer, H. Brunner, I. Grishkovskaya, K. Singh, A. Schleiffer, J. Jude, S. Deswal, R. Kalis, M. Vunjak, T. Lendl, R. Imre, E. Roitinger, T. Neumann, S. Kandolf, M. Schutzbier, K. Mechtler, G. A. Versteeg, D. Haselbach, J. Zuber, AKIRIN2 controls the nuclear import of proteasomes in vertebrates. Nature, 1–6 (2021).

41. H. Song, X. Feng, H. Zhang, Y. Luo, J. Huang, M. Lin, J. Jin, X. Ding, S. Wu, H. Huang, T. Yu, M. Zhang, H. Hong, S. Yao, Y. Zhao, Z. Zhang, METTL3 and ALKBH5 oppositely regulate m6A modification of TFEB mRNA, which dictates the fate of hypoxia/reoxygenation-treated cardiomyocytes. Autophagy. 15, 1419–1437 (2019).

42. D. Baillat, M.-A. Hakimi, A.M. Näär, A. Shilatifard, N. Cooch, R. Shiekhattar, Integrator, a Multiprotein Mediator of Small Nuclear RNA Processing, Associates with the C-Terminal Repeat of RNA Polymerase II. Cell. 123, 265– 276 (2005).

43. J. N. Jodoin, P. Sitaram, T. R. Albrecht, S. B. May, M. Shboul, E. Lee, B. Reversade, E. J. Wagner, L. A. Lee, Nuclear-localized Asunder regulates cytoplasmic dynein localization via its role in the integrator complex. Mol. Biol. Cell. 24, 2954–2965 (2013).

44. A. Malovannaya, Y. Li, Y. Bulynko, S. Y. Jung, Y. Wang, R. B. Lanz, B. W. O’Malley, J. Qin, Streamlined analysis schema for high-throughput identification of endogenous protein complexes. Proc. Natl. Acad. Sci. 107, 2431–2436 (2010).

45. K. Sabath, M. L. Stäubli, S. Marti, A. Leitner, M. Moes, S. Jonas, INTS10–INTS13–INTS14 form a functional module of Integrator that binds nucleic acids and the cleavage module. Nat. Commun. 11, 3422 (2020).

46. P. Lara-Gonzalez, J. Pines, A. Desai, Spindle assembly checkpoint activation and silencing at kinetochores. Semin. Cell Dev. Biol. 117, 86–98 (2021).

47. Y. J. Yang, A. E. Baltus, R. S. Mathew, E. A. Murphy, G. D. Evrony, D. M. Gonzalez, E. P. Wang, C. A. Marshall-Walker, B. J. Barry, J. Murn, A. Tatarakis, M. A. Mahajan, H. H. Samuels, Y. Shi, J. A. Golden, M. Mahajnah, R. Shenhav, C. A. Walsh, Microcephaly gene links trithorax and REST/NRSF to control neural stem cell proliferation and differentiation. Cell. 151, 1097–1112 (2012).

48. D. Jayaraman, B.-I. Bae, C. A. Walsh, The Genetics of Primary Microcephaly. Annu. Rev. Genomics Hum. Genet. 19, 177–200 (2018).

49. B. Neumann, T. Walter, J.-K. Hériché, J. Bulkescher, H. Erfle, C. Conrad, P. Rogers, I. Poser, M. Held, U. Liebel, C. Cetin, F. Sieckmann, G. Pau, R. Kabbe, A. Wünsche, V. Satagopam, M. H. A. Schmitz, C. Chapuis, D. W. Gerlich, R. Schneider, R. Eils, W. Huber, J.-M. Peters, A. A. Hyman, R. Durbin, R. Pepperkok, J. Ellenberg, Phenotypic profiling of the human genome by time-lapse microscopy reveals cell division genes. Nature. 464, 721–727 (2010).

50. G. Goshima, R. Wollman, S. S. Goodwin, N. Zhang, J. M. Scholey, R. D. Vale, N. Stuurman, Genes Required for Mitotic Spindle Assembly in Drosophila S2 Cells. Science. 316, 417–421 (2007).

51. I. M. Cheeseman, The kinetochore. Cold Spring Harb. Perspect. Biol. 6, a015826 (2014).

52. K. L. McKinley, I. M. Cheeseman, The molecular basis for centromere identity and function. Nat. Rev. Mol. Cell Biol. 17, 16–29 (2016).

53. M. Fischer, G.A. Müller, Cell cycle transcription control: DREAM/MuvB and RB-E2F complexes. Crit. Rev. Biochem. Mol. Biol. 52, 638–662 (2017).

54. T. Hayashi, Y. Fujita, O. Iwasaki, Y. Adachi, K. Takahashi, M. Yanagida, Mis16 and Mis18 Are Required for CENP-A Loading and Histone Deacetylation at Centromeres. Cell. 118, 715–729 (2004).

55. K. E. Gascoigne, K. Takeuchi, A. Suzuki, T. Hori, T. Fukagawa, I. M. Cheeseman, Induced Ectopic Kinetochore Assembly Bypasses the Requirement for CENP-A Nucleosomes. Cell. 145, 410–422 (2011).

56. K. L. McKinley, N. Sekulic, L. Y. Guo, T. Tsinman, B. E. Black, I. M. Cheeseman, The CENP-L-N Complex Forms a Critical Node in an Integrated Meshwork of Interactions at the Centromere-Kinetochore Interface. Mol. Cell. 60, 886–898 (2015).

57. J. van den Berg, A. G. Manjón, K. Kielbassa, F. M. Feringa, R. Freire, R. H. Medema, A limited number of double-strand DNA breaks is sufficient to delay cell cycle progression. Nucleic Acids Res. 46, 10132–10144 (2018).

58. J. P. Morgenstern, H. Land, Advanced mammalian gene transfer: high titre retroviral vectors with multiple drug selection markers and a complementary helper-free packaging cell line. Nucleic Acids Res. 18, 3587–3596 (1990).

59. D. Feldman, L. Funk, Pooled genetic perturbation screens with image-based phenotypes, OpticalPooledScreens. (2021), doi:https://doi.org/10.5281/zenodo.5002684.

60. S. Singh, M.-A. Bray, T. Jones, A. Carpenter, Pipeline for illumination correction of images for high-throughput microscopy. J. Microsc. 256, 231–236 (2014).

61. S. van der Walt, J. L. Schönberger, J. Nunez-Iglesias, F. Boulogne, J. D. Warner, N. Yager, E. Gouillart, T. Yu, scikit-image: image processing in Python. PeerJ. 2, e453 (2014).

62. J. Köster, S. Rahmann, Snakemake—a scalable bioinformatics workflow engine. Bioinformatics. 28, 2520– 2522 (2012).

63. F. Pedregosa, G. Varoquaux, A. Gramfort, V. Michel, B. Thirion, O. Grisel, M. Blondel, P. Prettenhofer, R. Weiss, V. Dubourg, J. Vanderplas, A. Passos, D. Cournapeau, M. Brucher, M. Perrot, É. Duchesnay, Scikit-learn: Machine Learning in Python. J. Mach. Learn. Res. 12, 2825–2830 (2011).

64. V. A. Traag, L. Waltman, N. J. van Eck, From Louvain to Leiden: guaranteeing well-connected communities. Sci. Rep. 9, 5233 (2019).

65. K. Jaqaman, D. Loerke, M. Mettlen, H. Kuwata, S. Grinstein, S. L. Schmid, G. Danuser, Robust single-particle tracking in live-cell time-lapse sequences. Nat. Methods. 5, 695–702 (2008).

66. J.-Y. Tinevez, N. Perry, J. Schindelin, G. M. Hoopes, G. D. Reynolds, E. Laplantine, S. Y. Bednarek, S. L. Shorte, K.W. Eliceiri, TrackMate: An open and extensible platform for single-particle tracking. Methods. 115, 80–90 (2017).

67. I. M. Cheeseman, A. Desai, A Combined Approach for the Localization and Tandem Affinity Purification of Protein Complexes from Metazoans. Sci. STKE. 2005, pl1–pl1 (2005).

68. J. C. Schmidt, H. Arthanari, A. Boeszoermenyi, N. M. Dashkevich, E. M. Wilson-Kubalek, N. Monnier, M. Markus, M. Oberer, R. A. Milligan, M. Bathe, G. Wagner, E. L. Grishchuk, I. M. Cheeseman, The Kinetochore-Bound Ska1 Complex Tracks Depolymerizing Microtubules and Binds to Curved Protofilaments. Dev. Cell. 23, 968–980 (2012).

69. J. Schindelin, I. Arganda-Carreras, E. Frise, V. Kaynig, M. Longair, T. Pietzsch, S. Preibisch, C. Rueden, S. Saalfeld, B. Schmid, J.-Y. Tinevez, D. J. White, V. Hartenstein, K. Eliceiri, P. Tomancak, A. Cardona, Fiji: an open-source platform for biological-image analysis. Nat. Methods. 9, 676– 682 (2012).

70. C. McQuin, A. Goodman, V. Chernyshev, L. Kamentsky, B. A. Cimini, K. W. Karhohs, M. Doan, L. Ding, S. M. Rafelski, D. Thirstrup, W. Wiegraebe, S. Singh, T. Becker, J. C. Caicedo, A. E. Carpenter, CellProfiler 3.0: Next-generation image processing for biology. PLOS Biol. 16, e2005970 (2018).

71. J. J. Trombetta, D. Gennert, D. Lu, R. Satija, A. K. Shalek, A. Regev, Curr. Protoc. Mol. Biol. 107, 4.22.1-17 (2014).

72. N. L. Bray, H. Pimentel, P. Melsted, L. Pachter, Near-optimal probabilistic RNA-seq quantification. Nat. Biotechnol. 34, 525–527 (2016).

73. M. D. Robinson, D. J. McCarthy, G. K. Smyth, edgeR: a Bioconductor package for differential expression analysis of digital gene expression data. Bioinformatics. 26, 139–140 (2010).

